# Spatiotemporal manipulation of ciliary glutamylation reveals its roles in intraciliary trafficking and Hedgehog signaling

**DOI:** 10.1101/282103

**Authors:** Shi-Rong Hong, Cuei-Ling Wang, Yao-Shen Huang, Yu-Chen Chang, Ya-Chu Chang, Ganesh V. Pusapati, Chun-Yu Lin, Ning Hsu, Hsiao-Chi Cheng, Yueh-Chen Chiang, Wei-En Huang, Nathan C. Shaner, Rajat Rohatgi, Takanari Inoue, Yu-Chun Lin

## Abstract

Tubulin post-translational modifications (PTMs) occur spatiotemporally throughout cells and are suggested to be involved in a wide range of cellular activities. However, the complexity and dynamic distribution of tubulin PTMs within cells have hindered the understanding of their physiological roles in specific subcellular compartments. Here we develop a method to rapidly deplete tubulin glutamlyation inside the primary cilia, a microtubule-based sensory organelle protruding on the cell surface, by targeting an engineered deglutamylase to the cilia in minutes. This rapid deglutamylation quickly leads to altered ciliary functions such as kinesin-2-mediated anterograde intraflagellar transport and Hedgehog signaling, along with no apparent crosstalk to other PTMs such as acetylation and detyrosination. Our study offers a feasible approach to spatiotemporally manipulate tubulin PTMs in living cells. Future expansion of the repertoire of actuators that regulate PTMs may facilitate a comprehensive understanding of how diverse tubulin PTMs encode ciliary as well as cellular functions.

## Introduction

The primary cilium is a microtubule-based sensory organelle protruding from the apical surface of resting cells; it is crucial in phototransduction, olfaction, hearing, embryonic development, and several cellular signaling pathways, such as Hedgehog (Hh) signaling^1, 2^. Defects in primary cilia lead to a number of human diseases^3^. Structurally, the cilium is composed of nine microtubule doublets called the axoneme, which offer mechanical support to the cilium, and also provide tracks for motor protein-dependent trafficking, known as intraflagellar transport (IFT)^4^. Polyglutamylation generates glutamate chains of varying lengths at the C-terminal tails of axonemal tubulin^5, 6^. This post-translational modification (PTM) occurs on the surface of microtubules and provides interacting sites for cellular components, such as microtubule-associated proteins (MAPs) and molecular motors^6^. However, the detailed mechanisms of how axonemal polyglutamylation regulates the stability and functionality of cilia remain to be understood.

Polyglutamylation is reversible, and tightly regulated by a balance between opposing enzymes for glutamylation or deglutamylation^7, 8^. More specifically, tubulin glutamylation is conducted by a family of tubulin tyrosine ligase-like (TTLL) proteins, including TTLL1, 4, 5, 6, 7, 9, 11, and 13^9, 10^. Each TTLL has a priority for initiation or elongation of glutamylation, as well as substrate preference between α- and β-tubulins^10^. This TTLL-mediated polyglutamylation is counteracted by a family of cytosolic carboxypeptidases (CCPs). Thus far, CCP1, 2, 3, 4, 5, and 6 have been identified as deglutamylases^6, 11^. CCP5 preferentially removes a glutamate at the branching fork, whereas other CCP members target a glutamate residue in a linear, tandem sequence in vivo^12, 13^. In contrast, Berezniuk *et al*. recently performed a biochemical assay to demonstrate that CCP5 cleaves glutamates at both locations and could complete the deglutamylation without the need for other CCP members^14^.

The effects of tubulin polyglutamylation on the structure and functions of microtubules have been studied mainly through the following approaches: 1) biochemical characterization of glutamylated microtubules, 2) cell biology assays for hyper- or hypo-glutamylation induced by genetically controlling the expression level of corresponding PTM enzymes, and 3) cell biological analysis of genetically mutated tubulins. As a result, it has been shown that chemical conjugation of glutamate side chains on purified microtubules increases the processivity and velocity of Kinesin-2 motors^15^. Tubulin hyperglutamylation leads to microtubule disassembly owing to the binding of a severing enzyme, namely spastin, to hyperglutamylated microtubules^16, 17^. Mice lacking a subunit of the polyglutamylase complex display hypoglutamylation in neuronal cells, which is accompanied by a decreased binding affinity of kinesin-3 motors to microtubules^18^. Moreover, the genetic or mopholino-mediated perturbation of polyglutamylases or deglutamylases across different model organisms results in morphological and/or functional defects in cilia and flagella^19–33^. Collectively, these studies strongly suggest the importance of tubulin polyglutamylation in the structural integrity and functionality of microtubules in cilia as well as other subcellular compartments. However, these approaches also revealed technical limitations. First, the distribution pattern of polyglutamylated tubulin is spatiotemporally dynamic; i.e., polyglutamylation is abundant in axoneme, centrioles, and neuronal axons in quiescent cells, which converges to the mitotic spindle and midbody during cytokinesis^6^. This dynamic feature cannot be directly addressed by conventional genetic manipulations or pharmacological inhibitors. Second, constitutive genetic perturbation often allows for compensation where cells adapt to their new genetic environment, likely leading to a missed detection of immediate consequences of loss-of- function such as an effect on transient interactions between tubulins with specific PTMs and their molecular partners^34^. Third, besides tubulin, many nucleocytoplasmic shuttling proteins such as nucleosome assembly proteins also have been identified as substrates of glutamylases^10, 35^. Therefore, global manipulation of genes encoding enzymes that modulate glutamylation is insufficient to specifically perturb axonemal glutamylation.

To circumvent these limitations, we developed a method named STRIP for <u>S</u>patio<u>T</u>emporal <u>R</u>ewriting of Intraciliary <u>P</u>TMs. The method is based on chemically inducible dimerization (CID) to spatiotemporally recruit a catalytic domain of deglutamylase onto ciliary axoneme, where the axonemal polyglutamylation can be rapidly stripped in an inducible manner. By implementing STRIP with simultaneous live-cell, time-lapse fluorescence imaging, we report cell biological analysis on the immediate consequences of deglutamylation that is rapidly induced inside primary cilia.

## Results

### Rapid relocalization of soluble proteins to the axoneme

We begin by describing a new approach, STRIP, for the spatiotemporal perturbation of axonemal polyglutamylation to study its immediate effect on the interplay among the microtubules, molecular motors, and MAPs (Fig. 1a). The basis of our technology is chemically inducible dimerization^36, 37^, in which a chemical such as rapamycin induces the dimerization of FK506 binding protein (FKBP) and FKBP-rapamycin binding domain (FRB) (Fig. 1a). To anchor a fusion protein of Cerulean3 and FRB onto ciliary axoneme, we employed microtubule-associated protein 4 (MAP4) that accumulates at the axoneme^38^. In particular, we used the truncated mutant of MAP4m containing partial proline-rich domain and affinity domain^38^ to minimize unexpected, yet possible biological effects arising from the N-terminal region (Supplementary Fig. 1a). When expressed in ciliated cells, Cerulean3-FRB-MAP4m mainly localized inside cilia and displayed weak signals in cytosol (Supplementary Fig. 1b). Its ciliary localization was further confirmed by not only overlapping distribution with a ciliary membrane marker, Arl13B, but also colocalization with two axoneme markers, acetylated tubulin and glutamylated tubulin (Supplementary Fig. 1b), assuring the intended localization of the MAP4m fusion protein. We then assessed the potential effect of overexpression of Cerulean3-FRB-MAP4m on the primary cilia. First, the expression of Cerulean3-FRB-MAP4m was confirmed to exhibit no significant effect on cilium length, axonemal acetylation, or axonemal glutamylation (Supplementary Fig. 1c). Next, we tested whether Cerulean3-FRB-MAP4m would affect IFT by monitoring Neon-IFT88, which labels IFT particles, with live-cell, time-lapse fluorescence microscopy^39^. The rates of IFT in anterograde and retrograde directions in the cilia with Cerulean3-FRB-MAP4m were not significantly different from those under control conditions (Supplementary Fig. 1d,e). Collectively, these results confirmed that Cerulean3-FRB-MAP4m is a valid fusion construct to be anchored at the ciliary axoneme.

**Figure 1.**
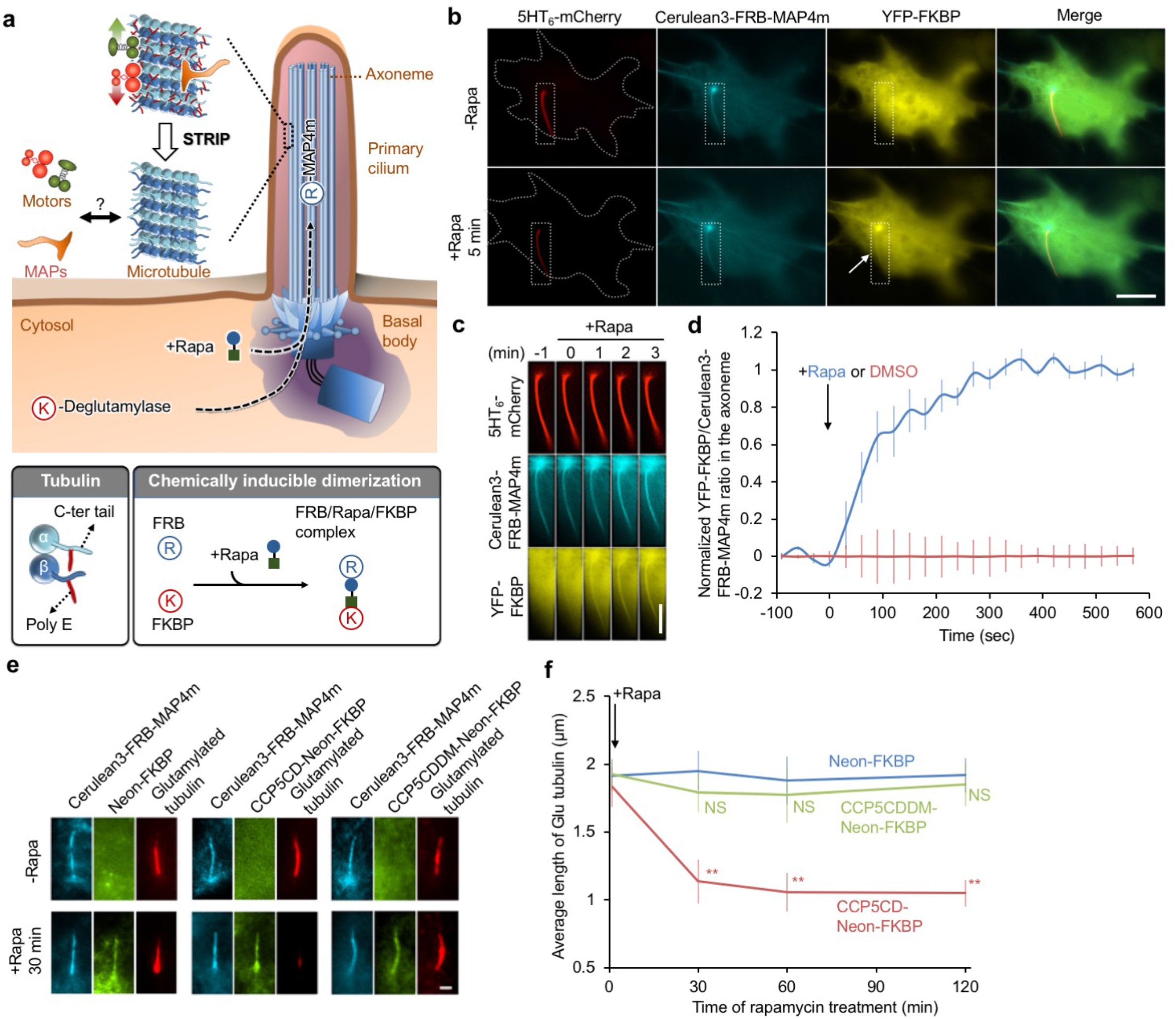
Spatiotemporally depleting tubulin glutamylation inside cilia. (**a**) Schematic diagram of the STRIP (<u>S</u>patio<u>T</u>emporal <u>R</u>ewriting of Intraciliary <u>P</u>TMs) approach which translocates an engineered deglutamylase from cytosol onto ciliary axonemes using the chemically induced dimerization system. The cytosolic FKBP-deglutamylase can be recruited to the FRB-MAP4m-labeled axoneme by adding rapamycin (+Rapa). The causal relationship among the axonemal glutamylation, motors and MAPs (microtubule-associated proteins) will be evaluated. (**b**) The addition of 100 nM rapamycin induces the accumulation of YFP-FKBP onto the Cerulean3-FRB-MAP4m-labeled axoneme (arrow) in NIH3T3 cells. Transfected cells at 80∼90% confluency were serum starved for 24 h and then incubated with 100 nM rapamycin for the indicated times. Scale bar, 10 μm. (**c**) Individual video frames in the axoneme region of cell in **b** upon 100 nM rapamycin treatment. Scale bar, 5 μm. Also, see Supplementary Movie 1. (**d**) Time course of YFP fluorescence intensity in the axoneme of NIH3T3 cells treated with 100 nM rapamycin (blue) or 0.1% DMSO (red). Data represent the mean ± s.e.m. (n = 10 cells for DMSO group, n = 8 cells for rapamycin group; four independent experiments). (**e,f**) Translocation of CCP5CD-Neon-FKBP to the axoneme reduces axonemal glutamylation. NIH3T3 cells were transfected with P2A-based constructs for co-expression of Cerulean3-FRB-MAP4m and Neon-FKBP-tagged proteins. Transfected cells at 80∼90% confluency were serum starved for 24 h and then incubated with 100 nM rapamycin for the indicated times. The level of axonemal glutamylation in cells expressing the indicated proteins was assessed by labeling with anti-glutamylated tubulin antibody and quantifying before and after the addition of 100 nM rapamycin. Scale bar, 1 μm. Data represent the mean ± s.e.m. (n = 276, 299, 244 cells for the Neon-FKBP, CCP5CD-Neon-FKBP, and CCP5CDDM-Neon-FKBP groups, respectively; 3∼5 independent experiments). NS and ** indicate no significant difference and *P* < 0.01, respectively, sbetween the control (Neon-FKBP) and the indicated groups.

Our previous study showing that most cytosolic proteins have access to the ciliary lumen^40^ suggests that cytosolic FKBP proteins can be trapped at the axoneme upon rapamycin addition in cells expressing Cerulean3-FRB-MAP4m. To test this possibility, we transfected NIH3T3 cells with Cerulean3-FRB-MAP4m (axoneme), 5HT_6_-mCherry (cilia membrane), and YFP-tagged FKBP (YFP-FKBP; cytosol). Subsequent exposure to rapamycin increased the YFP fluorescence signal at the ciliary axoneme over 5 min (Fig. 1b-d and Supplementary Movie 1). The time required for half-maximal accumulation of YFP-FKBP in the axoneme (t_1/2_) was 98.1 ± 24.0 s (mean ± s.e.m.; Fig. 1d). To generalize the method, we next tested the MAP4m approach with a gibberellin-based CID that works orthogonally to the rapamycin-based CID^41^. Here, we constructed two fusion proteins: a codon-optimized Gibberellin insensitivity DWARF1 (mGID1) fused to Neon (Neon-mGID1), and MAP4m fused to an N-terminal 92 amino acids of Gibberellin insensitive (GAIs-CFP-MAP4m). The addition of a gibberellin analog (GA_3_-AM) to cells co-expressing these fusion proteins led to accumulation of the Neon fluorescence signal at the ciliary axoneme (Supplementary Fig. 2 and Supplementary Movie 2), albeit with slower kinetics (127.3 ± 25.9 s) compared with the rapamycin-based CID (Supplementary Fig. 2c). Taken together, these results confirmed that the CID systems utilizing MAP4m can relocate cytosolic proteins onto the ciliary axoneme of living cells within minutes.

**Figure 2.**
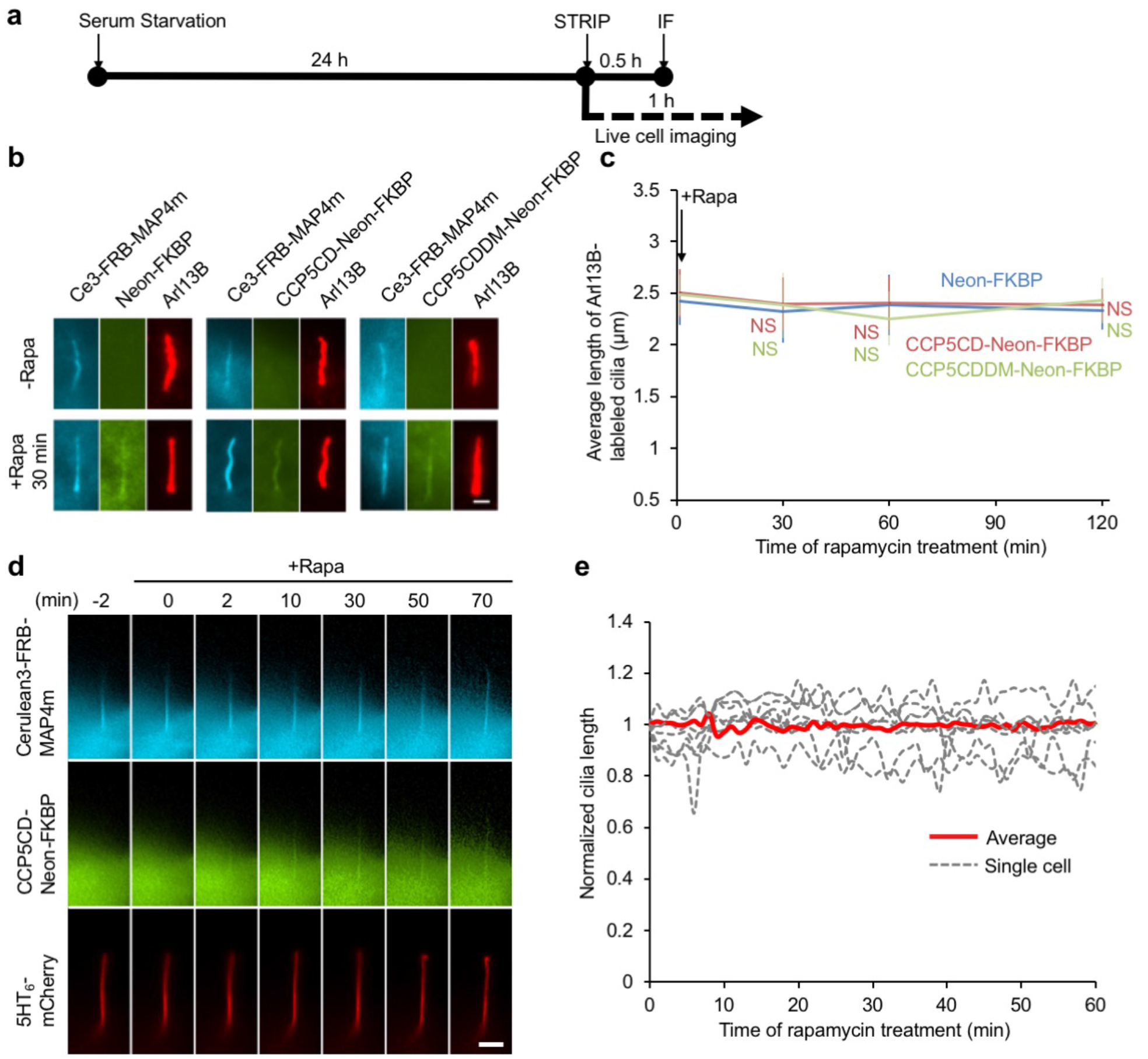
Glutamylation is not resquired for cilia maintenance. (**a**) The experimental procedure for applying STRIP together with immunofluorescence staining (IF) in **b** or live-cell imaging in **d** after serum starvation–induced ciliogenesis. (**b,c**) Rapid deglutamylation does not affect ciliary length in the steady state. NIH3T3 cells were transfected with P2A-based constructs for co-expression of Cerulean3 (Ce3)-FRB-MAP4m and Neon-FKBP-tagged proteins. Transfected cells at 80∼90% confluency were serum starved for 24 h and then incubated with 100 nM rapamycin for the indicated times. The cilium length in cells expressing the indicated proteins was measured by labeling with anti-Arl13B antibody and quantifying before and after the addition of 100 nM rapamycin. Scale bar, 1 μm. Data represent the mean ± s.e.m. (n = 269, 282, 238 cells for the Neon-FKBP, CCP5CD-Neon-FKBP, and CCP5CDDM-Neon-FKBP groups, respectively; 3∼5 independent experiments). NS indicates no significant difference between the control (Neon-FKBP) and the indicated groups. (**d**) The real-time morphology of primary cilium during axonemal deglutamylation. NIH3T3 cells were co-transfected with CCP5CD-Neon-FKBP-P2A-Cerulean3-FRB-MAP4m and a ciliary membrane marker, 5HT_6_-mCherry. Transfected cells at 80∼90% confluency were serum starved for 24 h and then incubated with 100 nM rapamycin for the indicated times. Shown are video frames of a cell co-expressing the indicated proteins upon treatment with 100 nM rapamycin. Scale bar, 4 μm. Also see Supplementary Movie 4. (**e**) The length of 5HT_6_-mCherry-labeled cilia in **d** was normalized by dividing the measured length by the length measured before rapamycin treatment and plotted. Red curves and gray dot curves show the average length and the length of each cilium during deglutamylation, respectively (n = 7 cilia from four independent experiments).

### Relocalizing CCP5 to the axoneme for rapid deglutamylation

To change the polyglutamylation status of tubulins in cilia, we implemented STRIP to recruit a deglutamylase to the axoneme. Among the tubulin deglutamylase family, CCP5 is demonstrated to efficiently remove glutamate residues at a branch point as well as those in a linear tandem both *in vitro* and *in vivo*^12, 14^, suggesting CCP5 as a valid candidate to actuate deglutamylation upon the STRIP execution. We thus constructed Neon-FKBP-tagged full-length CCP5 (CCP5FL-Neon-FKBP), intending to relocalizing this fusion protein from cytosol to axoneme upon rapamycin addition. However, CCP5FL-Neon-FKBP localized inside cilia even prior to rapamycin addition, thereby constitutively depleting axonemal glutamylation (Supplementary Fig. 3a-c), clearly hampering the use of the full-length CCP5 for the STRIP operation. Therefore, we assessed subcellular localization of truncation mutants of CCP5, namely N-terminus (CCP5N, residues 1-160) and catalytic domain (CCP5CD, residues 161-531). When expressed in cells as a fusion protein of Neon-FKBP, these two CCP5 truncations both localize in cytosol (Supplementary Fig. 3b). For further characterization of these truncation mutants, we introduced two point mutations (H252S and E255Q) to CCP5CD (CCP5CDDM) to impair the deglutamylation activity^9^. A fusion construct of this mutant (CCP5CDDM-Neon-FKBP) was also confirmed to be cytosolic (Supplementary Fig. 3b). In contrast to full-length CCP5, none of the truncated mutants impacted axonemal glutamylation (Supplementary Fig. 3b,c). To directly assess enzymatic activities of these CCP5 truncations, we forcefully anchored each of them to axoneme via fusion with MAP4m (Supplementary Fig. 4a). The immunofluorescence assay showed that CCP5CD, but not CCP5N or CCP5CDDM, significantly reduces axonemal glutamylation, indicating that CCP5CD retains viable enzyme activity (Supplementary Fig. 4b,c). Importantly, axonemal deglutamylation induced by CCP5FL-Neon-FKBP or CCP5CD-Cerulean3-MAP4m did not affect axonemal acetylation or cilium length (Supplementary Figs. 3c and 4c). Taken together, our results identified CCP5CD as an ideal candidate that becomes functional only at the axoneme without affecting other aspects of primary cilia such as cilia length and tubulin acetylation status.

**Figure 3.**
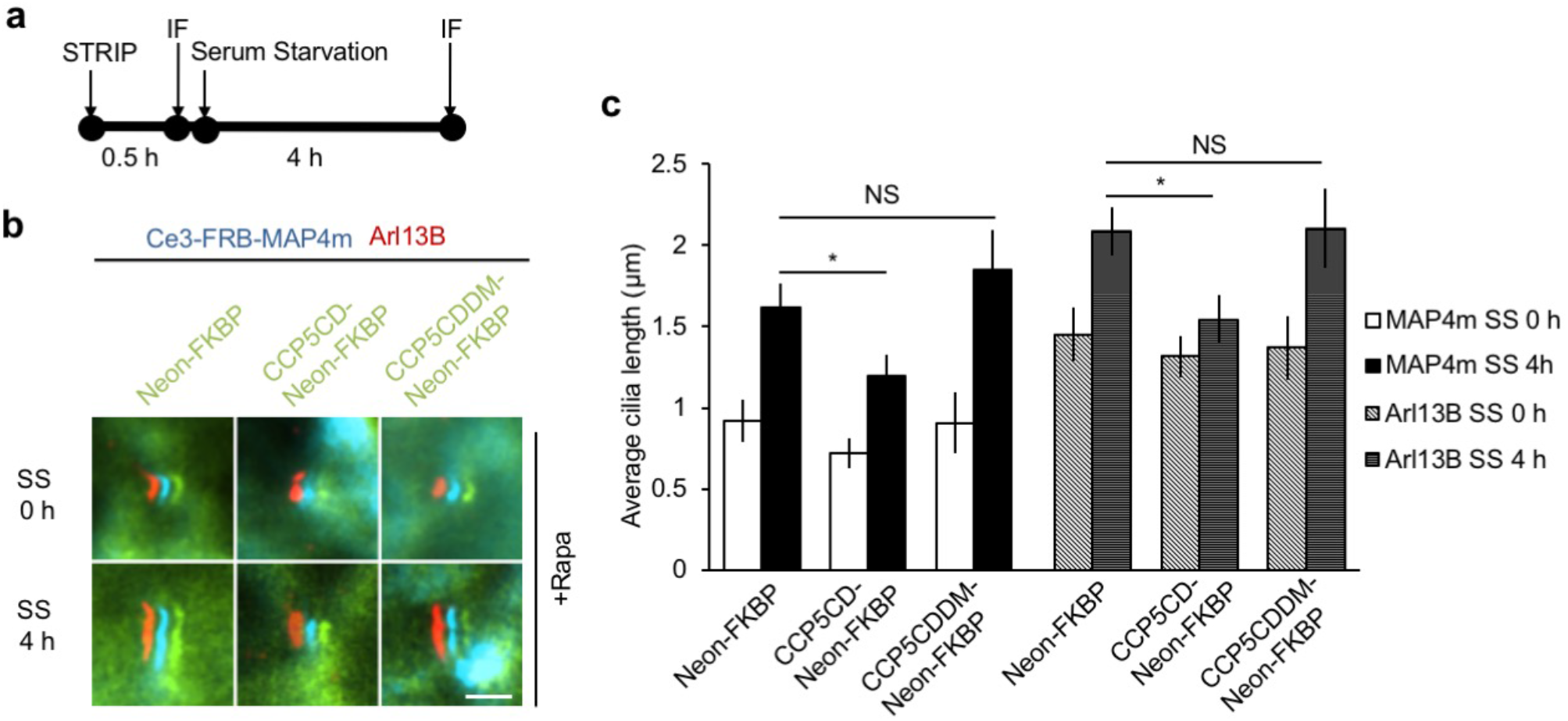
Glutamylation is important for cilia elongation. (**a**) Experimental procedure for applying STRIP together with immunofluorescence staining (IF) in **b** before and after serum starvation–induced cilia elongation. Washout of rapamycin after a 30-min incubation did not affect protein dimerization in cells owing to the irreversible nature of the CID system. (**b,c**) Rapid deglutamylation inhibits serum starvation–induced cilia elongation. NIH3T3 cells were transfected with P2A-based constructs for co-expression of Cerulean3-FRB-MAP4m and Neon-FKBP-tagged proteins. Transfected cells at 80∼90% confluency were incubated with 100 nM rapamycin for 30 min and then serum starved for the indicated times. The length of axoneme and of cilia was quantified in cells expressing the indicated proteins with rapamycin treatment before and after serum starvation (SS) for 4 h. The images show shifted overlays of the indicated proteins in cilia. Scale bar, 2 μm. (**c**) Average length of cilia labeled by MAP4m and Arl13B in **b**. Data represent the mean ± s.e.m. (n = 83, 80, 58, 59, 102, 92, 65, 82, 52, 51, 53, 51 cells from left to right; three independent experiments). NS and * indicate no significant difference and *P* < 0.05, respectively, between the control (Neon-FKBP) and the indicated groups.

**Figure 4.**
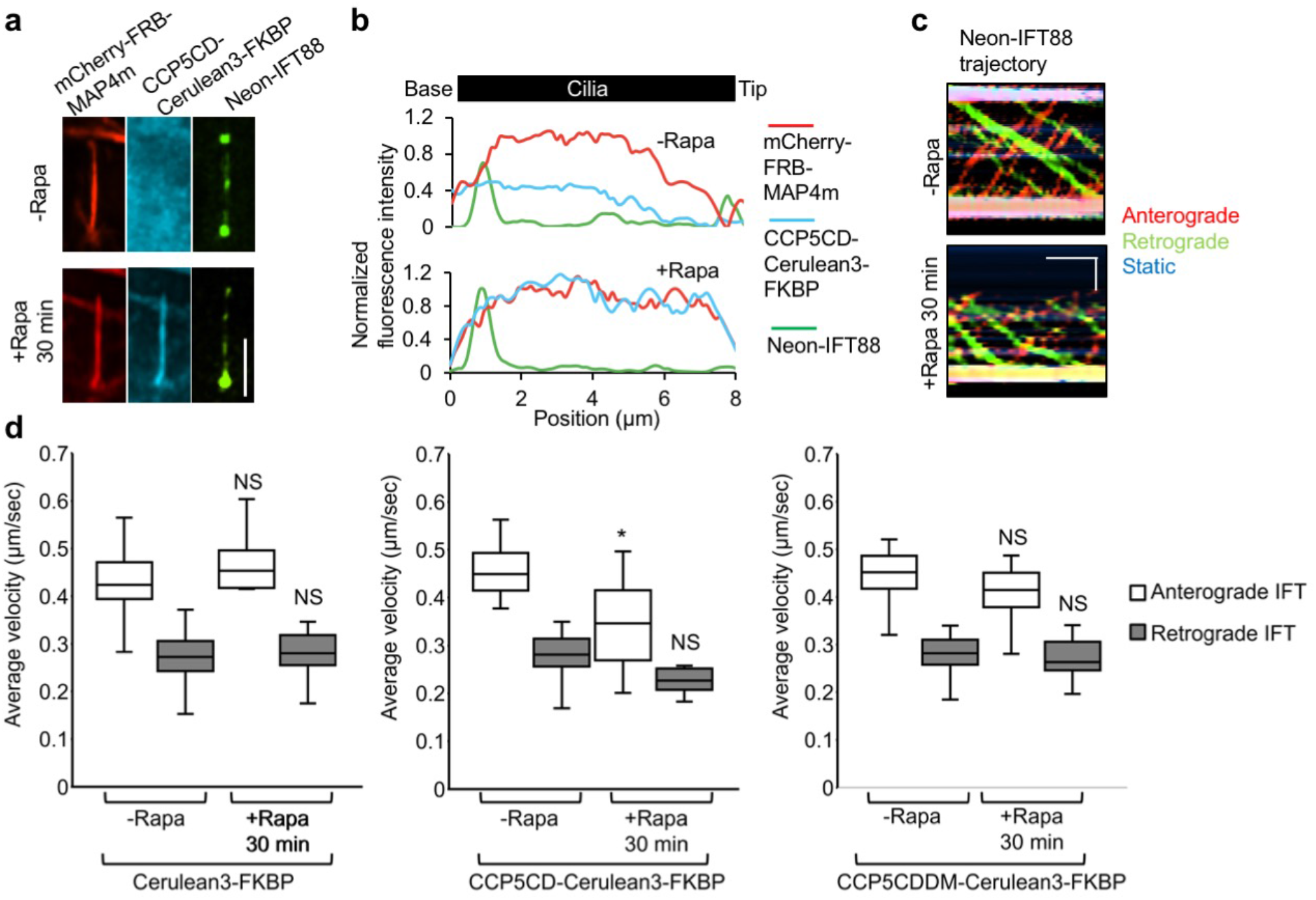
Rapid axonemal deglutamylation preferentially hampers anterograde IFT. (**a**) Neon-IFT88 accumulates at the base of deglutamylated cilia. Neon-IFT88 stable NIH3T3 cells were transfected with CCP5CD-Cerulean3-FKBP-P2A-mCherry-FRB-MAP4m. Transfected cells at 80∼90% confluency were serum starved for 24 h and then treated with 100 nM rapamycin for 30 min. Scale bar, 4 μm. (**b**) Linescan profile of the indicated proteins from base to the tip of primary cilium in **a**. (**c**) Representative kymographs of Neon-IFT88 generated from time-lapse imaging of cilium before and after rapamycin treatment for 30 min. Red, green, and blue lines represent the trajectories of Neon-IFT88 particles in anterograde and retrograde directions and static Neon-IFT88, respectively. Horizontal scale bar, 10 s. Vertical scale bar, 2 μm. Also, see Supplementary Movie 6. (**d**) Translocation of CCP5CD-Cerulean3-FKBP, but not Cerulean3-FKBP or CCP5CDDM-Cerulean3-FKBP, onto the axoneme hampers the IFT only in the anterograde direction. Neon-IFT88 stable NIH3T3 cells were transfected with P2A-based constructs for co-expression of mCherry-FRB-MAP4m and Cerulean3-FKBP-tagged proteins. Transfected cells at 80∼90% confluency were serum starved for 24 h and then treated with 100 nM rapamycin for the indicated times. The velocity of Neon-IFT88 was quantified according to the trajectories shown in the kymographs (see the method; n = 175, 150, 170 Neon-IFT88 particles for the Cerulean3-FKBP, CCP5CD-Cerulean3-FKBP, and CCP5CDDM-Cerulean3-FKBP groups, respectively; three independent experiments). NS and * indicate no significant difference and *P* < 0.05, respectively, between the conditions in the presence or absence of rapamycin.

To trigger deglutamylation of axonemal tubulin, CCP5CD-Neon-FKBP was recruited to the axoneme with Cerulean3-FRB-MAP4m by adding rapamycin (Supplementary Fig. 5 and Supplementary Movie 3). We inserted a viral P2A self-cleaving linker in between FKBP and FRB pieces, which encode relatively equal amounts of two proteins in subsequent experiments^42^. The kinetics of CCP5CD-Neon-FKBP accumulation in the axoneme upon rapamycin treatment were comparable to those of YFP-FKBP (CCP5CD-Neon-FKBP: 95.7 ± 21.9 s; YFP-FKBP: 98.1 ± 24.0 s). The recruitment of CCP5CD-Neon-FKBP to the axoneme reduced the glutamylation within 30 min (Fig. 1e,f). Interestingly, deglutamylation did not alter the localization of MAP4m, suggesting that MAP4m binding to microtubules is independent of tubulin glutamylation (Fig. 1e). Importantly, the recruitment of Neon-FKBP alone or catalytically inactive CCP5CDDM onto the axoneme was insufficient to deplete axonemal glutamylation, confirming that CCP5CD-induced axonemal deglutamylation is dependent on the enzymatic activity of CCP5 (Fig. 1e,f). It is noteworthy that the CCP5- or STRIP-induced deglutamylation left residual glutamylation at the proximal end of primary cilia (Fig. 1e and Supplementary Fig. 3b). The residual glutamylation localized at the inversin zone instead of the transition zone (Supplementary Fig. 6). The detailed mechanism of how axonemal tubulins in the inversin zone resist deglutamylation treatment is unclear.

**Figure 5.**
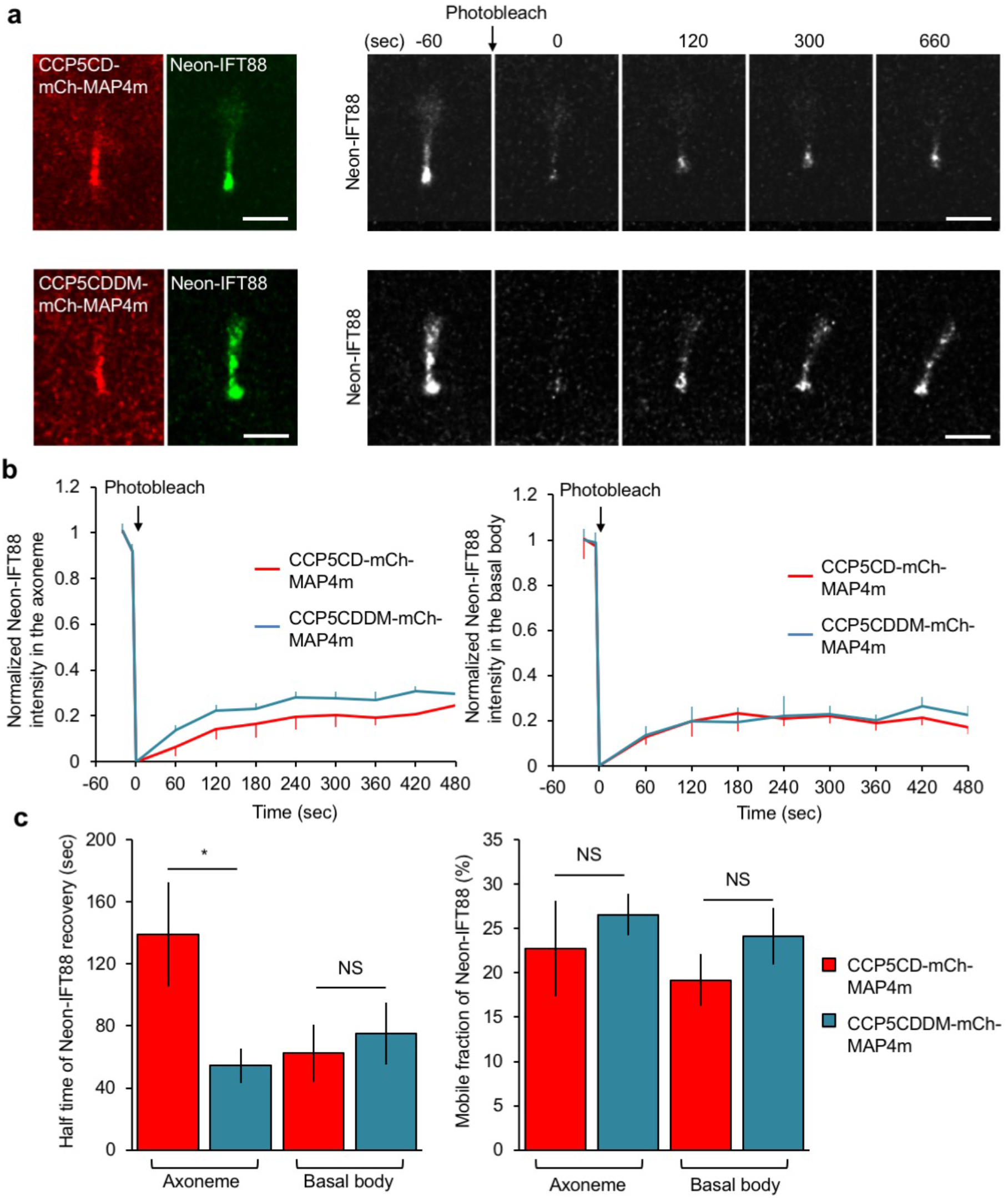
Axonemal deglutamylation hampers IFT dynamics along the axoneme but not the tethering of IFT machinery on the basal body. (**a**) Neon-IFT88 stable NIH3T3 cells were transfected with CCP5CD-mCherry-MAP4m or catalytically inactive CCP5CDDM-mCherry-MAP4m. Transfected cells at 80∼90% confluency were serum starved for 24 h prior to FRAP experiments. Cells expressing the indicated proteins were photobleached in the entire cilia region and allowed to recover for the indicated times. Scale bar, 2 μm. (**b**) Fluorescence recovery of Neon-IFT88 in the axoneme region (left) and basal body (right) was measured and plotted. (**c**) The recovery rate (left) and mobile fraction (right) of Neon-IFT88 in the axoneme and basal body were measured and plotted. Data represent the mean ± s.e.m. (n = 9, and 13 cilia for the CCP5CD-mCherry-MAP4m and CCP5CDDM-mCherry-MAP4m groups, respectively; four independent experiments).

**Figure 6.**
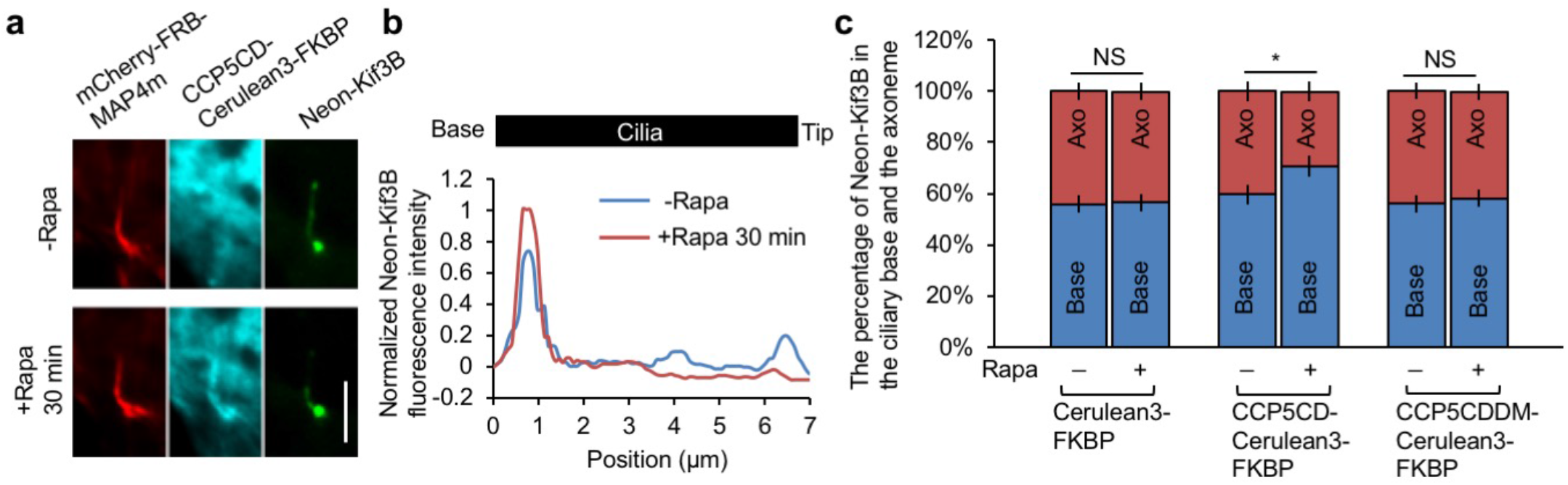
Kinesin 2 accumulates at the base of deglutamylated cilia. (**a**) Neon-Kif3B stable NIH3T3 cells were transfected with CCP5CD-Cerulean3-FKBP-P2A-mCherry-FRB-MAP4m. Transfected cells at 80∼90% confluency were serum starved for 24 h and then treated with 100 nM rapamycin for 30 min. Scale bar, 4 μm. (**f**) Linescan profile of Neon-Kif3B from the base to the tip of the cilium in **a**. (**c**) Neon-Kif3B stable NIH3T3 cells were transfected with P2A-based constructs for co-expression of mcherry-FRB-MAP4m and Cerulean3-FKBP-tagged proteins. Transfected cells at 80∼90% confluency were serum starved for 24 h and then treated with or without 100 nM rapamycin for 30 min. The percentage of Neon-Kif3B at the base (Base) or axoneme region (Axo) of control or deglutamylated cilia is plotted. Data represent the mean ± s.e.m. (n = 74, 48, 69 cells for the Cerulean3-FKBP, CCP5CD-Cerulean3-FKBP, and CCP5CDDM-Cerulean3-FKBP groups, respectively; three independent experiments). NS and * indicate no significant difference and *P* < 0.05, respectively.

### Specificity of CCP5-STRIP in quality and space

The status of polyglutamylation was verified by additional two antibodies, anti-polyE and anti-Δ2-tubulin antibodies, that recognize long glutamate side chains and penultimate glutamate in primary sequence of tubulin, respectively^12^. The recruitment of CCP5CD onto the axoneme does not significantly alter the level of long glutamate side chains (Supplementary Fig. 7a,b) or Δ2-tubulin (Supplementary Fig. 7c,d), suggesting the insufficient activity of CCP5 in shortening poly-glutamate side chains or catalyzing formation of Δ2-Δ3 tubulins, respectively (Supplementary Fig. 7a-d).

**Figure 7.**
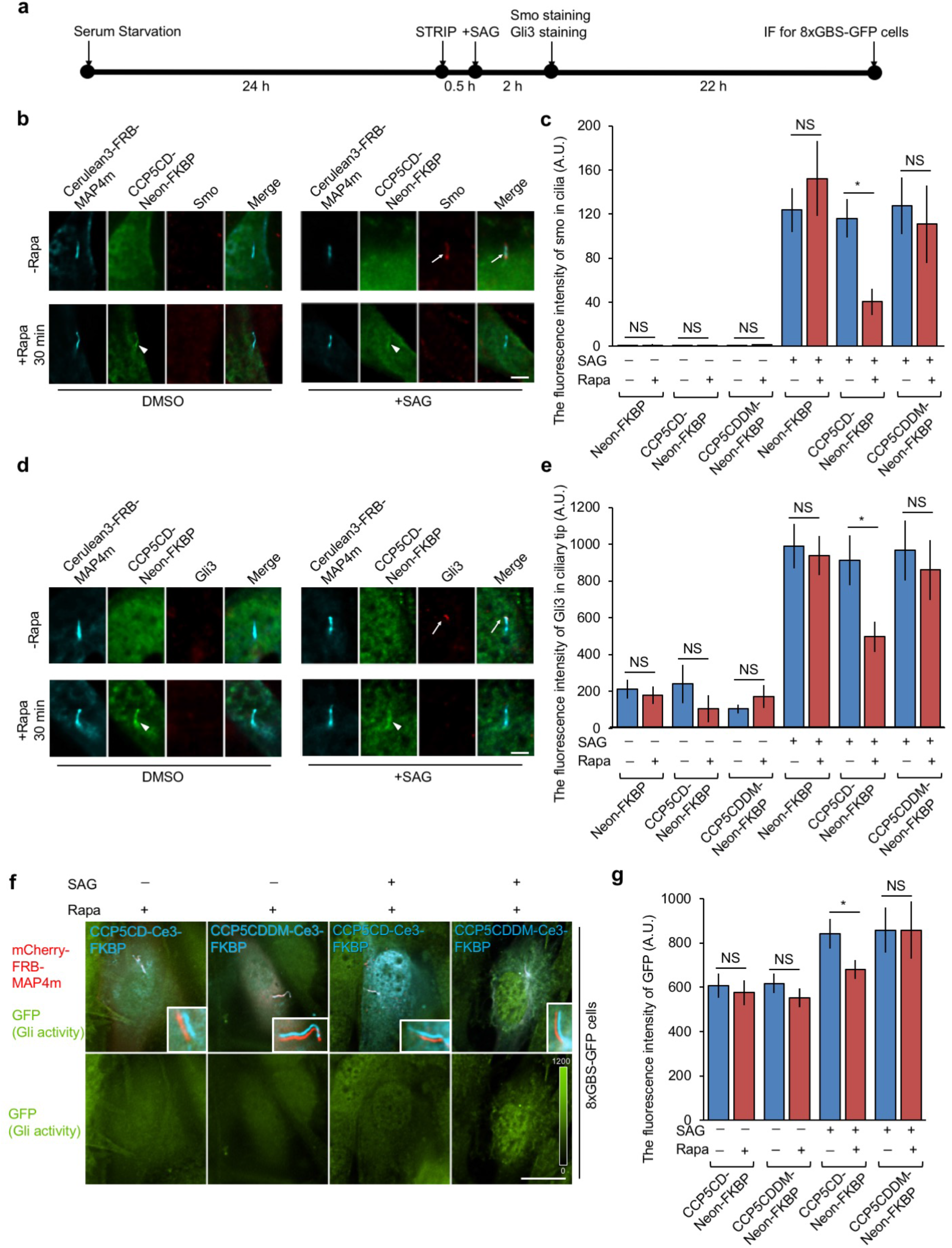
Axonemal deglutamylation suppresses Hedgehog signaling. (**a**) The experimental procedure for STRIP and Hedgehog induction. Washout of rapamycin after 30 min STRIP operation did not affect protein dimerization in cells owing to the irreversible nature of the CID system. (**b**) Axonemal deglutamylation reduces the level of Smoothened (Smo) in cilia after treatment with SAG. NIH3T3 cells were transfected with P2A-based constructs for co-expression of Cerulean3-FRB-MAP4m and Neon-FKBP-tagged proteins. Transfected cells at 80∼90% confluency were serum starved for 24 h and then incubated with 100 nM rapamycin or 0.1% DMSO for 30 min and subsequently treated with 1 μM SAG for 2 h. Scale bar, 5 μm. (**c**) Smoothened in cilia was quantified after the indicated treatments (n ≥ 18 cells from three independent experiments). (**d**) Axonemal deglutamylation reduces the level of Gli3 at the ciliary tip after treatment with SAG. NIH3T3 cells were transfected with P2A-based constructs for co-expression of Cerulean3-FRB-MAP4m and Neon-FKBP-tagged proteins. Transfected cells at 80∼90% confluency were serum starved for 24 h and then incubated with 100 nM rapamycin or 0.1% DMSO for 30 min and subsequently treated with 1 μM SAG for 2 h. Scale bar, 3 μm. (**e**) Gli3 at the ciliary tip was quantified after the indicated treatments. Data represent the mean ± s.e.m. (n ≥ 21 cells from three independent experiments). (**f**) Axonemal deglutamylation inhibits Gli activation upon Hedgehog stimulation. NIH3T3:8xGBS-GFP cells were transfected with P2A-based constructs for co-expression of mCherry-FRB-MAP4m and Cerulean3 (Ce3)-FKBP-tagged proteins. Transfected cells at 80∼90% confluency were serum starved for 24 h and then incubated with 100 nM rapamycin or 0.1% DMSO for 30 min and subsequently treated with 200 nM SAG for 24 h. The GFP intensity of cells was scaled to the same ranges in each image. The insets show the shifted overlays of the indicated proteins in cilia. Scale bar, 20 μm. **(g)** GFP intensity of NIH3T3:8xGBS-GFP was measured under the indicated conditions (n ≥ 13 cells from three independent experiments). Data represent the mean ± s.e.m. NS and * indicate no significant difference and *P* < 0.05, respectively.

To determine the spatial specificity of STRIP-triggered axonemal deglutamylation, we measured the level of tubulin glutamylation in other subcellular compartments. Glutamylated tubulins in the cytosol and basal body can be visualized after “cold treatment” before fixation (see the method, Supplementary Fig. 8a)^10^. Consistent with previous reports^8, 12^, deglutamylation by CCP5FL is prevalent in the cytosol, basal body, and axoneme (Supplementary Fig. 8a,b). In contrast, CCP5CD does not lead to noticeable change in tubulin glutamylation compared with a control condition with Neon alone. Strikingly, the STRIP-triggered deglutamylation was most prominent at the axonemes, and non-significant elsewhere in the same single cells, validating the high spatial precision (Supplementary Fig. 8a,b). It is somewhat interesting that expression of soluble CCP5CD in cells had only marginal effect on glutamylation in the cytosol, despite a nature of free diffusion. We speculate that the expression level of cytosolic CCP5CD may be low enough to take an effect, and/or that endogenous polyglutamylases in the cytosol may counteract CCP5CD-mediated deglutamylation in the cytosol. Nevertheless, a design principle of the STRIP enables drastic concentration of CCP5CD on the axoneme at the expense of a fraction of the CCP5CD pool in the cytosol, notably due to >10,000-fold volume difference between cilia and cytosol.

**Figure 8.**
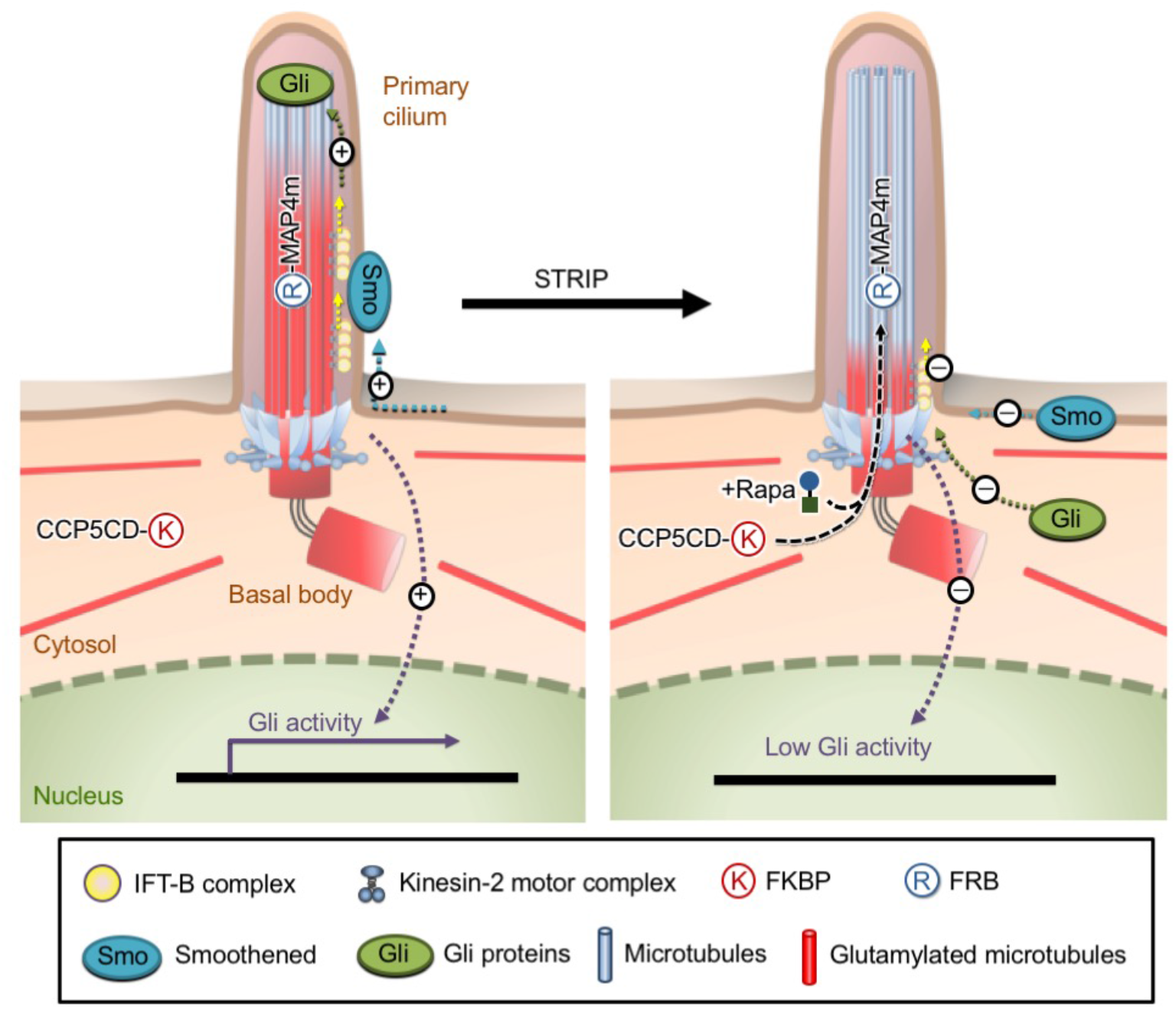
Model of our inducible axonemal deglutamylation system, STRIP, and proposed roles for tubulin glutamylation. The proximal end of axoneme is subject to polyglutamylation. Axonemal glutamylation facilitates kinesin-2-mediated anterograde IFT and the ciliary entry of the Hedgehog pathway components Smoothened (Smo) and Gli3 upon Hedgehog stimulation. The translocation of the CCP5 catalytic domain (CCP5CD) onto the MAP4m-appended axoneme by adding rapamycin specifically strips axonemal glutamylation and subsequently inhibits anterograde IFT and Hedgehog signaling without noticeably affecting ciliary structure in the steady state.

### The roles of glutamylation in ciliary structure

To reveal the effect of axonemal deglutamylation on the ciliary structure, we first measured the cilia length based on Arl13B staining. The cilium length measured before and after STRIP treatment at least for 2 h was comparable among the control (no CCP5), wild-type (CCP5CD), and inactive mutant (CCP5CDDM) (Fig. 2b,c). Next, we assessed length and morphology of primary cilia in live cells. Cells expressing CCP5CD-Neon-FKBP, Cerulean3-FRB-MAP4m, and 5HT_6_-mCherry underwent live-cell fluorescence imaging. Rapamycin addition does not result in alteration of the cilium length or morphology (Fig. 2d,e and Supplementary Movie 4). As expected from the result with CCP5CD-Cerulean3-MAP4m, STRIP-mediated deglutamylation does not affect axonemal detyrosination and acetylation (Supplementary Figs. 7e,f, 9). These results indicate that the structural integrity of steady-state primary cilia is organized independently of glutamylation.

Axoneme growth takes place in an early stage of ciliogenesis, and associates with polyglutamylation^43, 44^. To study the role of polyglutamylation in growing axoneme, we rapidly deglutamylated axonemes under a serum starved condition to induce cilia elongation (Fig. 3). MAP4m accumulates with no problem at the growing axoneme, enabling the STRIP operation for deglutamylation (Supplementary Fig. 10 and Supplementary Movie 5). Recruitment of CCP5CD onto the growing axoneme significantly slows the extension of cilium length (Fig. 3 b,c). Taken together, these results indicate that axonemal glutamylation is important for cilia elongation during ciliogenesis, but not for maintenance of steady-state cilia.

### Deglutamylation hampers kinesin-2-mediated anterograde IFT

Besides mechanical support, the axoneme also serves as railways for IFT^45, 46^. We thus evaluated whether axonemal deglutamylation impacts the rate of IFT. To test this, the IFT activity was quantified before and after STRIP-mediated axonemal deglutamylation by monitoring Neon-IFT88 with live-cell fluorescence microscopy (Fig. 4). A subsequent linescan analysis of the Neon-IFT88 signal in individual cilia at two time points (before and 30 min after rapamycin treatment) showed that Neon-IFT88 initially spread across the entire cilia in the form of puncta converges to the base of the cilium after deglutamylation (Fig. 4a,b). To assess IFT motility, a kymograph was then generated based on the time-lapse images of Neon-IFT88 before and after deglutamylation (Fig. 4c). The velocity of IFT in both directions (i.e., anterograde and retrograde) for control and deglutamylated cilia was calculated based on the slopes of each IFT88 trajectory on the kymograph (Fig. 4c). The rates of bidirectional IFT were comparable in both control groups (Fig. 4d, left and right panels). In contrast, anterograde but not retrograde IFT was significantly slowed after axonemal deglutamylation by CCD5CD (0.46 ± 0.03 μm/s vs. 0.37 ± 0.03 μm/s) (Fig. 3d, middle panel and Supplementary Movie 6). These results demonstrated that rapid deglutamylation preferentially affects anterograde IFT.

Besides rapid deglutamylation, we also evaluated the effect of long-term deglutamylation on IFT dynamics by measuring the velocity of Neon-IFT88 in cilia expressing CCP5CD-mCherry-MAP4m (Supplementary Fig. 11). Cilia expressing catalytically inactive CCP5CDDM-mCherry-MAP4m were included as a control (Supplementary Fig. 11). Consistent with the finding that rapid deglutamylation attenuated anterograde IFT, long-term deglutamylation induced by CCP5CD-mCherry-MAP4m significantly slowed anterograde IFT but not retrograde IFT (Supplementary Fig. 11b). Moreover, long-term deglutamylation only slightly but not significantly decreased the frequency of Neon-IFT88 in both directions (Supplementary Fig. 11c).

We next used fluorescence recovery after photobleaching (FRAP) to measure the ciliary entry of IFT in control and deglutamylated cilia (Fig. 5). Neon-IFT88 in the whole ciliary region was photobleached, and the fluorescence recovery at the basal body or axoneme was measured. The recovery rate and mobile fraction of Neon-IFT88 at the ciliary base were compared between control and deglutamylated cilia (Fig. 5 and Supplementary Movie 7), which indicated that axonemal deglutamylation does not affect the tethering of Neon-IFT88 to the basal body. Interestingly, axonemal deglutamylation induced by CCP5CD-mCherry-MAP4m significantly slowed the recovery rate of Neon-IFT88 onto the axoneme (t_1/2_ in control cilia: 54.43 ± 11.24 s; t_1/2_ in deglutamylated cilia: 139.14 ± 33.49 s), probably owing to defects in anterograde IFT induced by axonemal deglutamylation (Fig. 5 and Supplementary Movie 7). Taken together, these results confirmed that axonemal deglutamylation slows anterograde IFT without blocking the tethering of the IFT machinery to the basal body.

Anterograde IFT is mainly powered by kinesin motors^45^, whose distribution and motility are reportedly modulated by tubulin glutamylation^15, 18^. We therefore hypothesized that axonemal deglutamylation inhibits anterograde IFT by impairing kinesin motility. To address this, we examined the effects of axonemal deglutamylation on the distribution of Neon-Kif3B, a motor subunit of the kinesin-2 complex. Neon-Kif3B was observed to move along the axoneme (Supplementary Fig. 12a,b and Supplementary Movie 8). The anterograde rate of Neon-Kif3B particles was measured to be comparable to that of the Neon-IFT88 trains (Supplementary Fig. 12b,c), suggesting that Neon-Kif3B can be used to assess the motility of kinesin motors. We then found that Neon-Kif3B accumulates at the proximal end of deglutamylated cilia (Fig. 6a,b), just like Neon-IFT88 (Fig. 4a,b). To reinforce this finding, we performed quantification of Neon-Kif3B distribution in the cilia divided into two compartments, base and axoneme, which exhibited more Neon-Kif3B at the base of deglutamylated cilia compared with control cilia (70.7 ± 2.2% vs. 59.6 ± 3.9%) (Fig. 6c). In summary, these results suggest that glutamylation controls anterograde IFT through Kif3B of the kinesin-2 complex.

The Lechtreck group once showed that the demand of anterograde IFT in newly growing cilia is much higher than that in mature cilia^47^. Therefore, the rapid deglutamylation-induced defects in anterograde IFT may cause more severe inhibition of cilia elongation than cilia maintenance, a prediction consistent with our finding that rapid deglutamylation impaired elongation of immature, but not mature cilia (Figs. 2, 3).

### Axonemal deglutamylation inhibits Hh signaling

We next investigated whether axonemal deglutamylation affects ciliary Hh signaling. The Hh receptor Patched and the orphan G protein–coupled receptor (Gpr161) localize in cilia and block Hh signaling in the absence of Hh ligands. These two signaling molecules exit from the cilium once the Hh pathway is activated^48^. Immunofluorescence assay revealed that STRIP-induced axonemal deglutamylation did not affect the ciliary exit of Patched1-YFP and GPR161 upon the treatment of Smo agonist, SAG (Supplementary Fig. 13), indicating that glutamylation is not involved in modulating these two upstream signaling components in Hh signaling.

The kinesin-2 complex interacts with Hh components to regulate Hh signaling^48–52^, making us hypothesize that deglutamylation-induced defects in kinesin-2-mediated anterograde IFT would collaterally affect Hh signaling. We thus examined the subcellular location of Hh components such as Smoothened (Smo) and Gli3 in control and deglutamylated cilia after stimulation with the SAG (Fig. 7a). The SAG-induced accumulation of Smo (Fig. 7b,c) and Gli3 (Fig. 7d,e) in the cilia was significantly impaired by axonemal deglutamylation. The distribution and motility of GFP-Gli3 in deglutamylated cilia upon treatment with SAG was also evaluated by FRAP (Supplementary Fig. 14). Deglutamylation induced by CCP5CD-mCherry-MAP4m suppressed the ciliary distribution of GFP-Gli3 upon stimulation with SAG (Supplementary Fig. 14b), which is consistent with the results obtained with STRIP operation (Fig. 7d,e). Moreover, axonemal deglutamylation significantly slowed the entry of GFP-Gli3 into cilia, probably owing to defective anterograde IFT (Supplementary Fig. 14c,d and Supplementary Movie 9).

We also tested whether axonemal deglutamylation inhibits Gli activation using 8xGBS-GFP:NIH3T3 reporter cells in which a minimal promoter and 8xGli-Binding-Site (GBS) drive green fluorescent protein (GFP) expression^53^. GFP served as a fluorescence reporter to represent Gli transcription activities. As expected, treatment with SAG significantly increased GFP intensity in these reporter cells with no STRIP operation (Cells expressing CCP5CDDM-Cerulean3-FKBP; Fig. 7f,g). However, STRIP-mediated axonemal deglutamylation abolished the GFP fluorescence increase in SAG-treated cells (Cells expressing CCP5CD-Cerulean3-FKBP; Fig. 7f.g). Taken together, these results confirmed that axoneme glutamylation is important for Hh signaling, likely through an anterograde IFT-dependent mechanism.

## Discussion

With the newly developed STRIP technique, we stripped off polyglutamate modification of axonemal tubulins in living cells within minutes, and revealed that glutamylated tubulins positively regulate at least three characteristics of the primary cilia (Fig. 8): 1) elongation of nascent axonemes during ciliogenesis (Fig. 3), 2) anterograde IFT mediated by kinesin-2 (Figs. 4-6,8 and Supplementary Fig. 11), and 3) Hh signaling induced by Smo agonist (Figs. 7-8). In contrast, glutamylation played little role in other tubulin PTMs such as acetylation and detyrosination (Supplementary Figs. 7 and 9), retrograde IFT (Fig. 4 and Supplementary Fig. 11), or length maintenance of mature cilia (Fig. 2 and Supplementary Figs. 3, 4, 9).

Defective ciliogenesis was previously observed after complete genetic depletion or mutation of kinesin-2 in ciliated organisms^54–57^, which seems at first glance inconsistent with our present finding that deglutamylation-induced defects in kinesin-2 does not affect cilia maintenance. There are at least two explanations for this apparent discrepancy. In addition to axoneme, kinesin-2 also localizes at the ciliary base to regulate its organization^58^ (Fig. 6a and Supplementary Fig. 12a). Therefore, genetic manipulation of kinesin-2 may have affected functions of kinesin-2 not only at the axoneme but also the ciliary base, possibly resulting in defective basal bodies that could devastate ciliogenesis. Another explanation is that residual anterograde IFT in the STRIP cells may have been sufficient for cilia growth and maintenance, unlike the cases for near complete loss-of-function of kinesin-2 proteins^54–57^.

Our work showed that axonemal deglutamylation preferentially hampers anterograde IFT but not retrograde IFT (Fig. 4c.d and Supplementary Fig. 11). In several ciliated model organisms, glutamylation is primarily abundant on the B-tubules of the outer axoneme doublets^43, 59–61^. Polyglutamylase and deglutamylase mutations in several studies frequently cause defects of B-tubules in the axoneme doublets^20, 21, 62^, with one exception that abnormal A-tubules were observed in TTLL6 morphants^25^. Pigino and colleagues used correlative fluorescence and three-dimensional electron microscopy to elegantly demonstrate that anterograde IFT trains move along the B-tubule, while retrograde IFT uses the A-tubule in *Chlamydomonas* flagella^46^. Thus, it will be interesting to determine which tubule doublet (A, B or both) causes the defective anterograde IFT after tubulin deglutamylation. In addition to mammalian primary cilia and *Chlamydomonas* flagella, several recent studies demonstrated that perturbation of specific tubulin isotype and tubulin glutamylation affects ciliary ultrastructure and impacts the motility of different anterograde kinesin motors in C. *elegans* cilia^63, 64^. The relationship among tubulin codes, microtubule ultrastructure, and molecular motors in different ciliated model organisms merit comprehensive scrutiny.

By verifying the effect of deglutamylation on the distribution of various Hh signaling molecules as well as Gli transcriptional activities, we claimed that axonemal glutamylation positively regulates Hh signaling presumably through anterograde IFT-dependent mechanism (Fig. 7 and Supplementary Figs. 13 and 14). Our results showed that axonemal deglutamylation attenuates Hh signaling mainly by disturbing the ciliary entry of Smo and Gli proteins instead of the removal of their upstream negative regulators from cilia (Fig. 7 and Supplementary Figs. 13 and 14). Previous studies found that ciliary entry of Gli protein is driven by anterograde IFT, which offers a legitimate explanation on how deglutamylation-induced defects in anterograde IFT inhibit Hh signaling^49, 52^. However, several studies using pulse-chase assay and single molecule imaging have demonstrated that ciliary entry of Smo depends on lateral diffusion rather than anterograde IFT^65, 66^. Further work is required to decipher whether kinesin 2 assists Smo in crossing the diffusion barrier at cilia base^48, 67^.

One of the advantages of the STRIP is its modular nature originating from the CID approach^37^ with which CCP5 and MAP4m can be respectively replaced with other tubulin modification enzymes and other microtubule-binding proteins^68^. By implementing both rapamycin- and gibberellin-mediated STRIPs, it would be possible to rewrite two different PTMs. Moreover, our dimerization approach can be advanced to reversible operation by utilizing light-induced dimerization systems such as Cry2-CIBN and iLid-SspB^69^. Such an enhanced STRIP will enable targeting of a specific pair of tubulin PTMs and tubulin subtypes, which may help understand the tubulin code more thoroughly in the near future.

## Methods

### Cell culture and transfection

NIH3T3 cells, CFP-FRB-MAP4m-expressing stable NIH3T3 cells, Neon-IFT88-expressing stable NIH3T3 cells, and Neon-Kif3B-expressing stable NIH3T3 cells were maintained at 37 °C in 5% CO_2_ in DMEM (Corning) supplemented with 10% fetal bovine serum (Gibco), penicillin and streptomycin (Corning). NIH3T3:8xGBS-GFP reporter line was cultured in DMEM containing 10% FBS and 2 mM GlutaMAX (GIBCO). To induce ciliogenesis, cells were serum starved for 24 h. For induction of Hh pathway, ciliated cells were treated with 1 μM SAG (Enzo) for 2 h or 200 nM SAG for 24 h. Plasmid DNA transfection was performed by LT-1 transfection reagent (Mirus) or X-tremegene 9 (Roche) 24 h prior to the serum starvation. Transfected cells were treated with 100 nM rapamycin (LC Laboratories) or 100 μM GA_3_-AM^41^ for rapid induction of protein dimerization and translocation in living cells.

### Generation of NIH3T3 stable cell lines

NIH3T3 cells stably expressing respective proteins were generated using MSCV retroviral expressing system. The open reading frames of respective proteins together with CMV promoter were subcloned into MSCV vector (Clontech) by in-Fusion HD cloning kit (Clontech). Platinum 293T cells were transfected with MSCV-CMV-CFP-FRB-MAP4m, MSCV-CMV-Neon-IFT88, or MSCV-CMV-Neon-Kif3B for generating retrovirus particles. Retrovirus harboring respective genes was incubated with NIH3T3 cells in the presence of 10 μg/ml polybrene (Sigma) and infected cells were selected by 2.5 μg/ml puromycin (Sigma).

### Live cell-imaging

Cells were plated on poly(D-lysine)-coated borosilicate glass Lab-Tek 8-well chambers (Thermo Scientific). Live-cell imaging was performed using a Nikon T1 inverted fluorescence microscope (Nikon) with a 60× oil objective (Nikon), DS-Qi2 CMOS camera (Nikon) and 37°C, 5% CO_2_ heat stage (Live Cell Instrument). Imaging was acquired using Nikon element AR software. Images with multiple z-stacks were processed with Huygens Deconvolution (Scientific Volume Imaging) and the maximum intensity projections of images were produced by Nikon elements AR software (Nikon). The image analysis was mainly conducted by Nikon elements AR software (Nikon).

### Measurement of IFT dynamics

Neon-IFT88 or Neon-Kif3B stable NIH3T3 cells were transfected with indicated constructs and then seeded onto poly(D-lysine)-coated borosilicate glass Lab-Tek 8 well chambers (Thermo Scientific). The cells were imaged every 200 ms for 30 s on a Nikon T1 inverted fluorescence microscope (Nikon) with a 60× oil objective (Nikon), DS-Qi2 CMOS camera (Nikon) and 37°C, 5% CO_2_ heat stage (Live Cell Instrument). Time-lapse images were processed by Huygens deconvolution (Scientific Volume Imaging) and Kymographs were produced with Image J and the plugin KymographClear^70^.

### Immunofluorescence staining

Cells were plated on poly(D-lysine)-coated borosilicate glass Lab-Tek 8-well chambers (Thermo Scientific), fixed in 4% paraformaldehyde (Electron Microscopy Sciences) at room temperature for 10 min, then in 100% methanol (Sigma-Aldrich) at −20°C for 4 min. Fixed sample was permeabilized by 0.1% Triton X-100 (Sigma-Aldrich) and then incubated in blocking solution (PBS with 2% bovine serum albumin) for 30 min at room temperature. To label glutamylated tubulin in the cytosol and basal body, cells were cold treated at 4°C for 1 h and then fixed in cold methanol at −20°C for 10 min. Primary antibodies were diluted in blocking solution and were used to stain the cells for 1 h at room temperature. Primary antibodies used in this study were anti-glutamylated tubulin (1:500 dilution; AG-20B-0020, AdipoGen), anti-Gli3 (1:200 dilution; AF3690, R&D Systems), anti-acetylated tubulin (1:500 dilution, T7451, Sigma-Aldrich), anti-Smoothened (1:100 dilution, ab 38686, Abcam), anti-Arl13B (1:500 dilution, ab83879, Abcam), anti-polyglutamate chain (1:500 dilution, IN105, AdipoGen), anti-Δ2 tubulin (1:500 dilution, AB3203, Merck Millipore), anti-detyrosinated tubulin (1:500 dilution, AB3201, Millipore), and anti-GPR161 (1:200 dilution; 13398-1AP, Proteintech). Secondary antibodies were diluted in blocking solution (1:1000 dilution) and were incubated with sample for 1 h at room temperature.

### FRAP experiments

FRAP experiments were carried out with a Nikon A1 confocal system (Nikon) and 100× oil objective (Nikon). Before bleaching, two sequential images were taken to obtain a baseline of fluorescence intensity of Neon-IFT88 or GFP-Gli3. The cilia region of cells expressing GFP-Gli3 or Neon-IFT88 was then photobleached and allowed to recover for 20 min. The fluorescence intensity of GFP-Gli3 or Neon-IFT88 was measured with Nikon elements AR software (Nikon).

### Statistical analysis

Statistical analysis was performed with an unpaired two-tailed Student’s *t* test and whether variances were equal or not was determined by F test. *P* values were calculated, when *P* value ≥0.05 represent no significant difference, *P* value < 0.05 represent significant difference, and *P* value < 0.01 represent highly significant difference.

### Data availability

The data that support the findings of this study are available from the corresponding author upon reasonable request.

*Note: Supplementary information is available in the online version of the paper*

## Acknowledgements

We thank Dr. W. James Nelson (Stanford University) for the MAP4m construct, Dr. Gregory J. Pazour (University of Massachusetts Medical School) for the IFT88 and Kif3B constructs, Dr. Maarten F. Bijlsma and Dr. Helene Damhofer (University of Amsterdam) for the GBS-GFP construct, Dr. Koji Ikegami (Hamamatsu University School of Medicine) for CCP5 construct, and Dr. Jin-Wu Tsai (National Yang-Ming University) for the Patched1-YFP construct. We also thank Dr. Kristen Verhey (University of Michigan Medical School) for helpful discussions. We are grateful to Dr. Koji Ikegami (Hamamatsu University School of Medicine) for critical reading of the manuscript as well as Robert DeRose for writing suggestions. We thank Emily Su (Johns Hopkins University) for assistance with experiments and Dr. Tasuku Ueno (University of Tokyo) for the synthesis of GA_3_-AM. This study was supported in part by the National Institutes of Health (GM105448 and GM118082 to R.R.; R01DK102910 to T.I.), the Ministry of Science and Technology (MOST), Taiwan (MOST 104-2311-B-007-001, MOST 105-2628-B-007-001-MY3, and Program for Translational Innovation of Biopharmaceutical Development-Technology Supporting Platform Axis-nMACS imaging to Y.C.L.), and start-up funding from National Tsing Hua University to Y.C.L.

## Author contributions

S.R.H., C.L.W., Y.S.H., Y.C.C., and Y.C.L. generated DNA constructs, and S.R.H., C.L.W., Y.S.H., Y.C.L., Y.C.C., Y.C.C., C.Y.L., N.H., H.C.C., Y.J.C., and W.E.H. performed cell biology experiments and quantified the imaging results. Y.C.L. and C.L.W. performed live-cell imaging. N.S. generated the Neon plasmid. G.V.P. established the NIH3T3:8xGBS-GFP reporter line under the supervision of R.R. S.R.H., C.L.W., T.I., and Y.C.L. wrote the paper.

## Figure legends

**Supplementary Figure 1.**
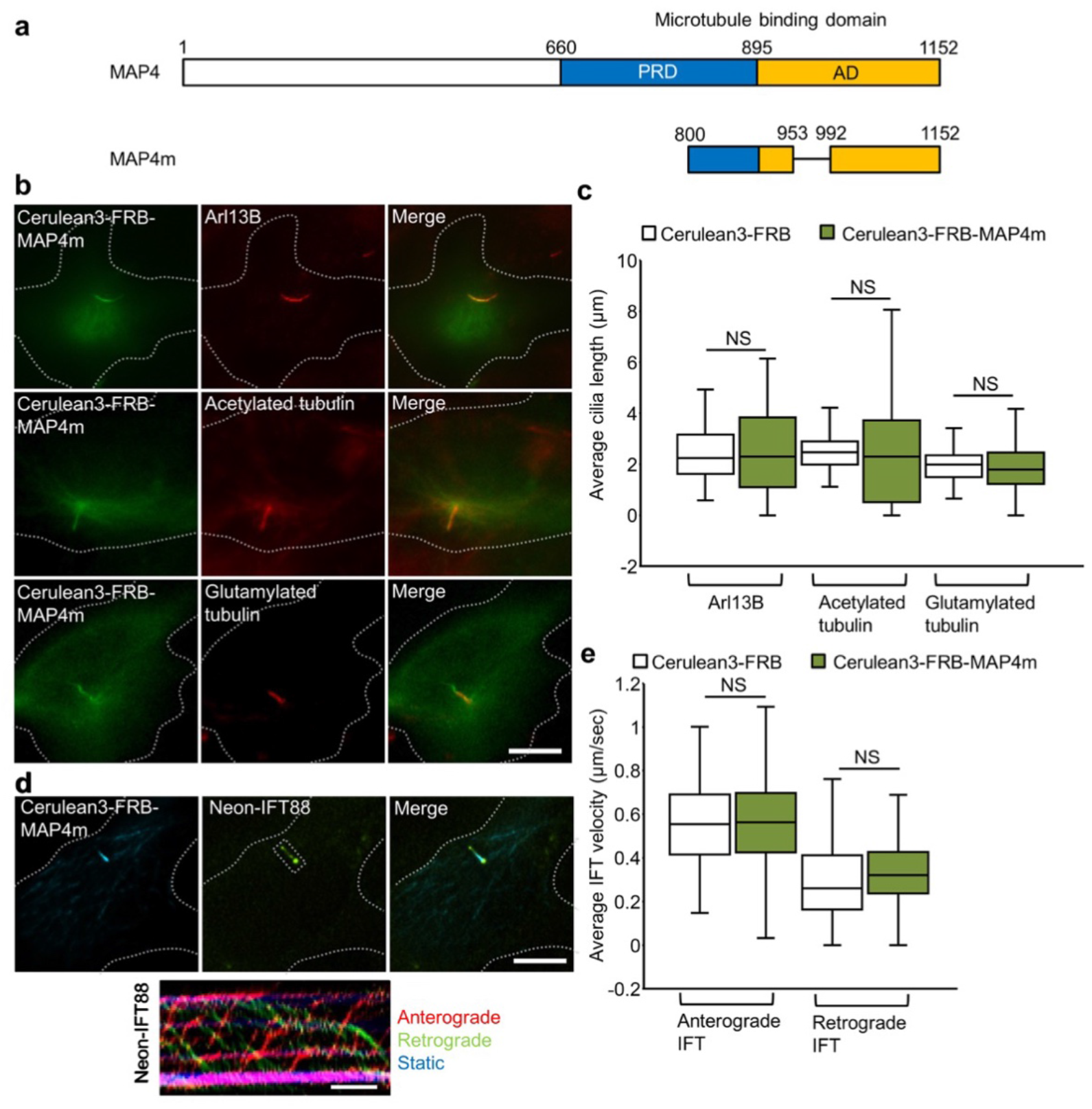
MAP4m localizes at the ciliary axoneme without affecting the ciliary structure and intraflagellar transport. (**a**) Schematic diagram of MAP4 and truncated MAP4m protein. The proline-rich domain (PRD) and affinity domain (AD) in MAP4 protein are shown. (**b**) Cerulean3-FRB-MAP4m (green) localizes at the axoneme in NIH3T3 cells. NIH3T3 cells were transfected with Cerulean3-FRB-MAP4m. Transfected cells at 80∼90% confluency were serum starved for 24 h and then immunostained by the indicated antibodies. Arl13B marks ciliary membrane (red). Acetylated tubulin and glutamylated tubulin mark the axoneme (red). Scale bars, 5 μm. (**c**) Expression of MAP4m does not affect cilium length. Box plots of cilium length measured using Arl13B, acetylated tubulin, glutamylated tubulin shown in **b**. (n = 69, 36, 36, 106, 35, 146 from left to right; three independent experiments) (**d**) Expression of MAP4m does not affect the dynamics of IFT. The Neon-IFT88 stable NIH3T3 cells were transfected with Cerulean3-FRB or Cerulean3-FRB-MAP4m. Transfected cells at 80∼90% confluency were serum starved for 24 h and imaged by time-lapse imaging. The kymograph of IFT dynamics was created by KymographClear. Scale bar, 10 s. (**e**) The rates of IFT were tracked and quantified by the movement of Neon-IFT88 in anterograde and retrograde directions. (n = 105, 90, 100, 66 Neon-IFT88 particles from left to right; 3-5 independent experiments). NS represents no significant difference between the control group and the cells expressing Cereulan3-FRB-MAP4m.

**Supplementary Figure 2.**
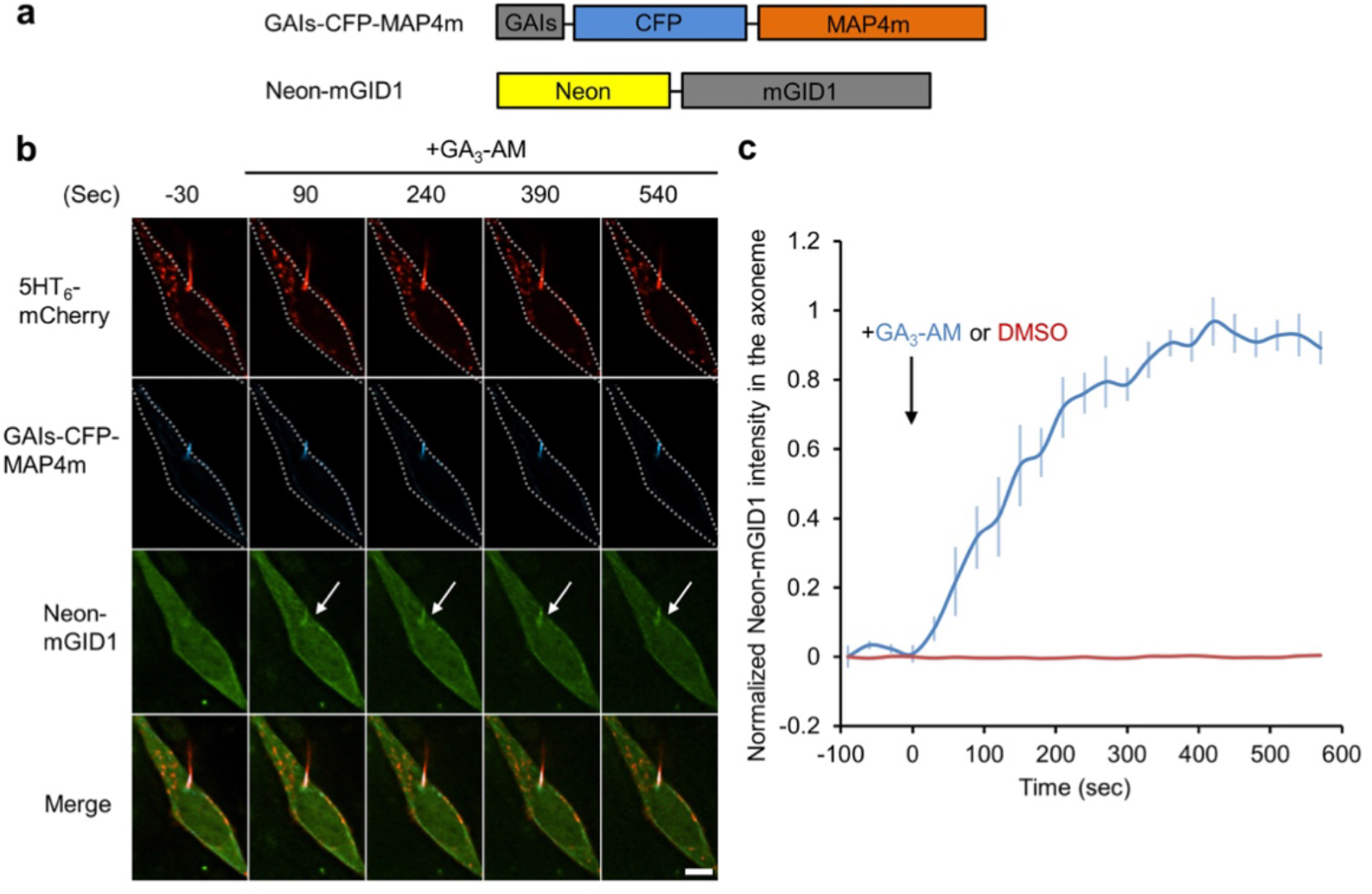
Translocation of cytosolic proteins onto the axoneme using a gibberellin-induced dimerization system. (**a**) Schematic diagram of the constructs that encoded GAIs-CFP-MAP4m and Neon-mGID1 proteins. (**b**) The addition of 100 μM GA_3_-AM induces accumulation of Neon-mGID1 onto the GAIs-CFP-MAP4m-labeled axoneme (arrows). NIH3T3 cells were co-transfected with 5HT_6_-mCherry, GAIs-CFP-MAP4m and Neon-mGID1. Transfected cells at 80∼90% confluency were serum starved for 24 h and then treated with 100 μM GA_3_-AM for the indicated time. 5HT_6_-mCherry marks ciliary membrane. Scale bar, 5 μm. Also, see Supplementary Movie 2. (**c**) Time course of Neon-mGID1 fluorescence intensity in the axoneme treated with 100 μM GA_3_-AM (blue) or 0.1% DMSO (red). (n = 6 cells in the GA_3_-AM treated group; n = 8 cells in the DMSO treated group; three independent exeriments) Data represent the mean ± s.e.m.

**Supplementary Figure 3.**
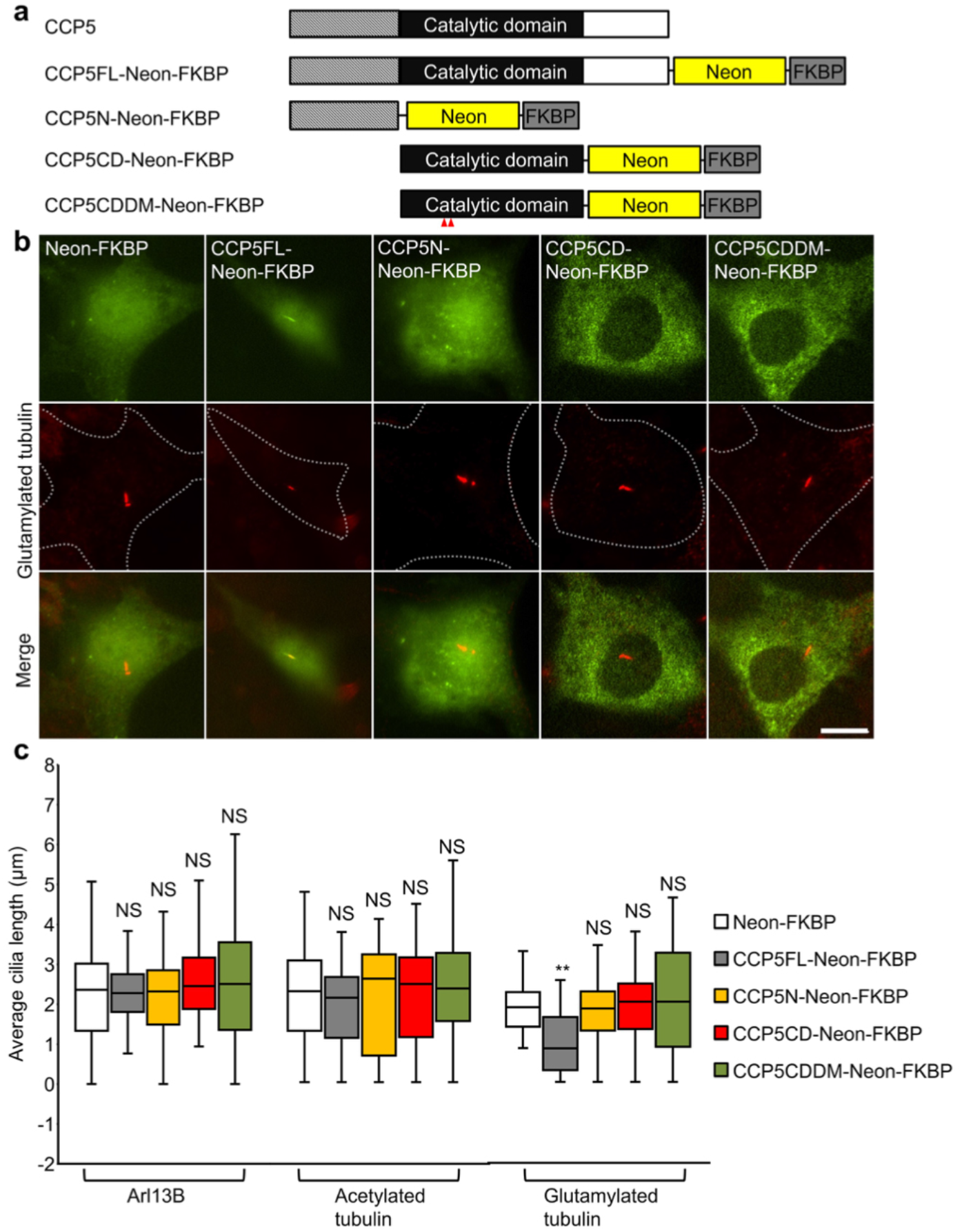
Characterization of various domains in CCP5 protein. (**a**) Schematic diagram of Neon-FKBP tagged full length (CCP5FL-Neon-FKBP), N-terminus (CCP5N-Neon-FKBP), wild type catalytic domain (CCP5CD-Neon-FKBP), and catalytically inactive form of CCP5 (CCP5CDDM-Neon-FKBP). (**b**) CCP5FL-Neon-FKBP but not CCP5N-Neon-FKBP, CCP5CD-Neon-FKBP, or CCP5CDDM-Neon-FKBP (green) localizes in cilia and reduces the axonemal glutamylation. NIH3T3 cells were transfected with the indicated Neon-FKBP-tagged proteins. Transfected cells at 80∼90% confluency were serum starved for 24 h, followed by immunostained with glutamylated tubulin (red). Scale bar, 10 μm. (**c**) CCP5FL-Neon-FKBP but not other CCP5 domain significantly reduces the axonemal glutamylation without affecting the ciliary length and axonemal acetylation. Box plots of cilia length labeled by antibodies against Arl13B, acetylated tubulin, and glutamylated tubulin, respectively, are shown. (n = 67, 30, 31, 45, 30, 44, 30, 33, 31, 47, 50, 47, 36, 33, 34 cells from left to right; 3-5 independent experiments). NS and ** represent no significant difference and *P* < 0.01 between the control (Neon-FKBP) and the cells expressing indicated proteins.

**Supplementary Figure 4.**
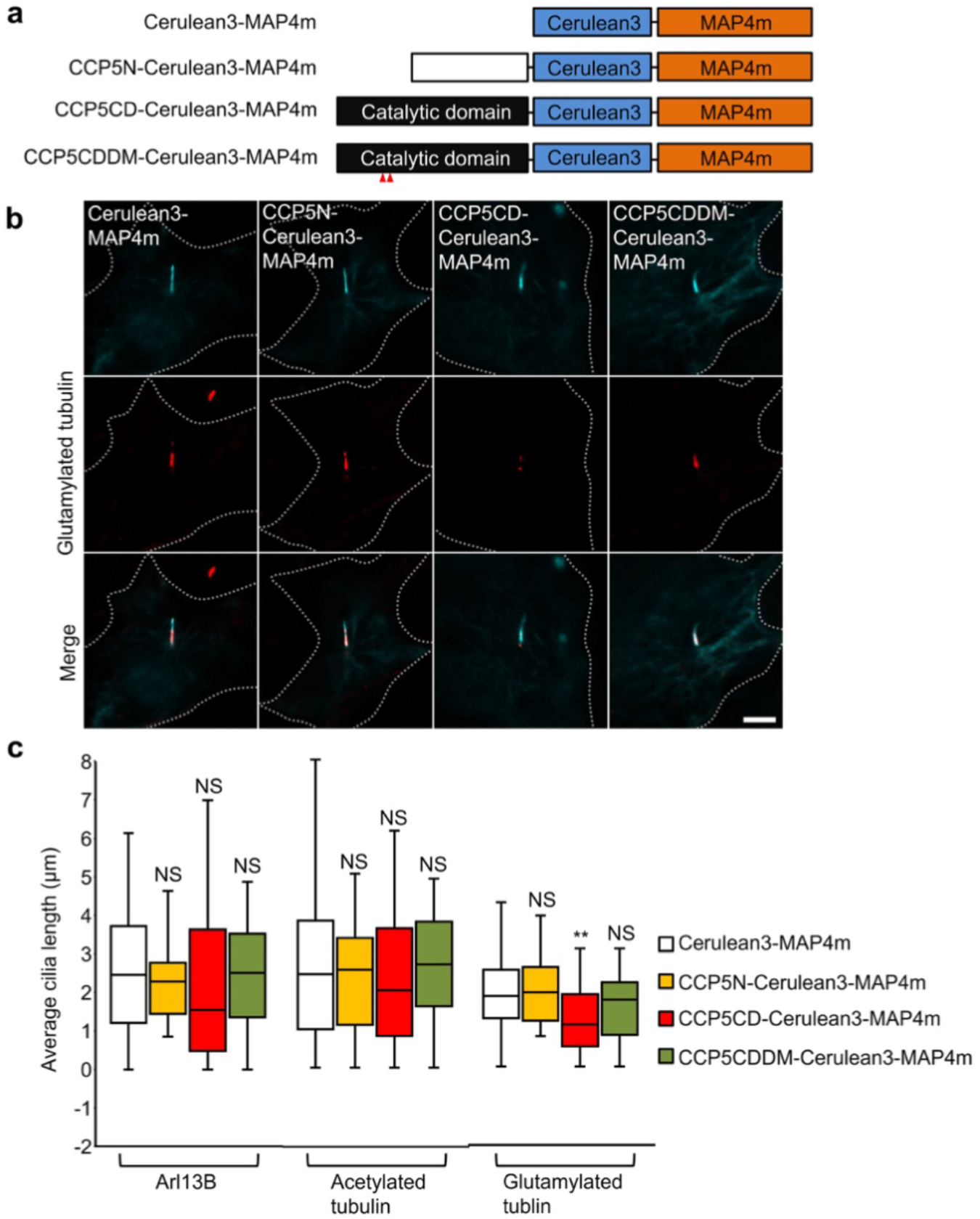
Targeting CCP5CD to the axoneme reduces axonemal glutamylation. (**a**) Schematic diagram of the various Cerulean3-MAP4m-fused CCP5N, CCP5CD, and CCP5CDDM. (**b**) CCP5CD-Cerulean3-MAP4m localizes in the axoneme and reduces axonemal glutamylation. NIH3T3 cells were transfected with indicated Cerulean3-MAP4m-tagged proteins. Transfected cells at 80∼90% confluency were serum starved for 24 h and immunostained for glutamylated tubulin (red). Scale bar, 5 μm. (**c**) CCP5CD-Cerulean3-MAP4m but not other CCP5 constructs significantly reduces the axonemal glutamylation without affecting ciliary length and axonemal acetylation. Box plots of cilia length labeled by antibodies against Arl13B, acetylated tubulin, and glutamylated tubulin, respectively, are shown. (n = 94, 50, 30, 36, 146, 60, 49, 30, 127, 31, 41, 34 cells from left to right; four independent experiments). NS and ** represent no significant difference and *P* < 0.01 between the control (Cerulean3-MAP4m) and the cells expressing indicated proteins.

**Supplementary Figure 5.**
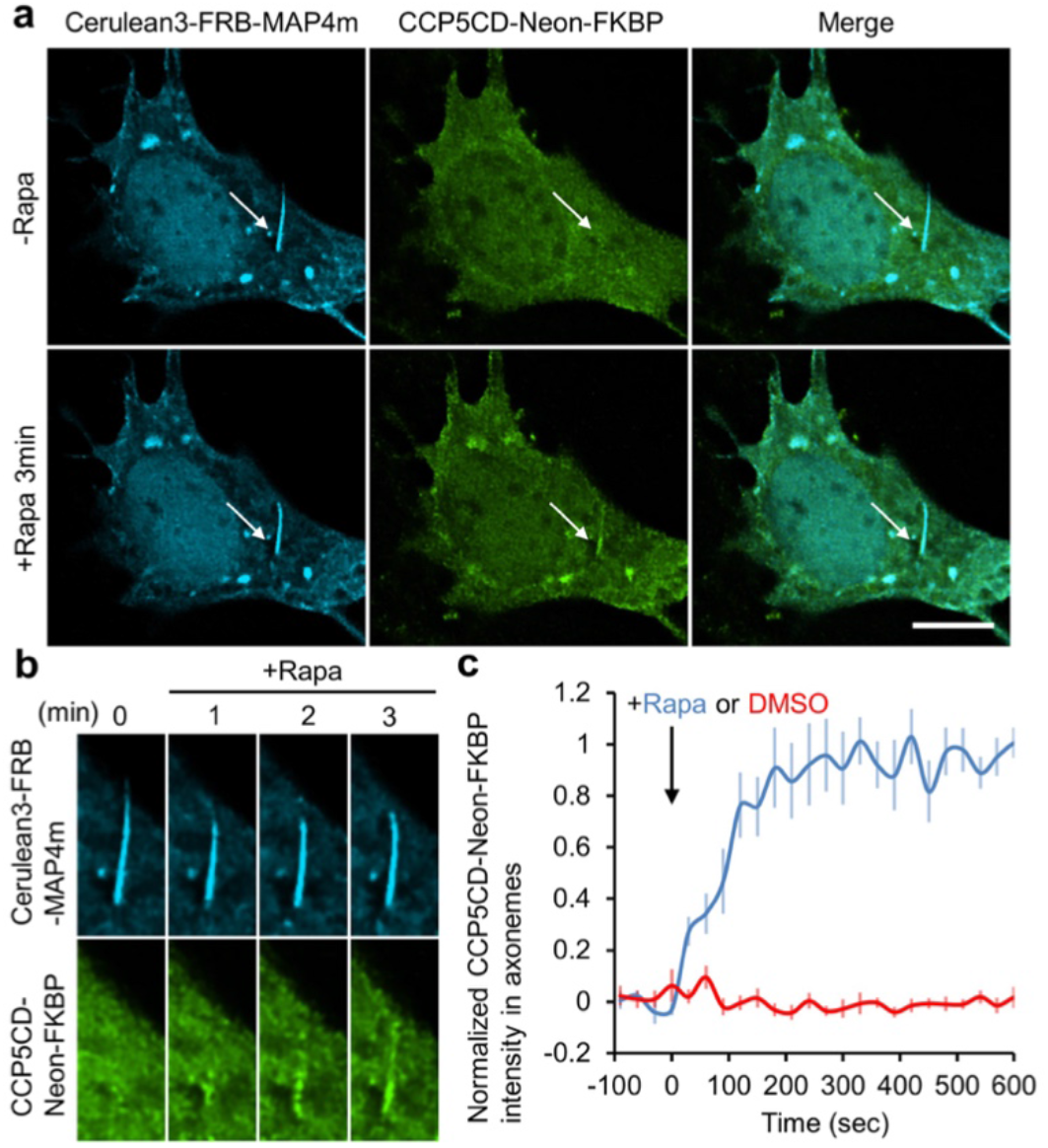
Translocation of CCP5CD-Neon-FKBP onto the ciliary axoneme using a rapamycin dimerization system. (**a**) The addition of 100 nM rapamycin induces accumulation of CCP5CD-Neon-FKBP onto the Cerulean3-FRB-MAP4m-labeled axoneme (arrows). NIH3T3 cells were co-transfected with Cerulean3-FRB-MAP4m and CCP5CD-Neon-FKBP. Transfected cells at 80∼90% confluency were serum starved for 24 h and treated with 100 nM rapamycin. Scale bar, 5 μm. Also, see Supplementary Movie 3. (b) Individual video frames in the axoneme region of cell in **a** upon 100 nM rapamycin treatment. (**d**) Time course of YFP fluorescence intensity in the axoneme of NIH3T3 cells treated with 100 nM rapamycin (blue) or 0.1% DMSO (red). Data represent the mean ± s.e.m. (n = 11 cells for the DMSO group, n = 7 cells for the rapamycin group; three independent experiments).

**Supplementary Figure 6.**
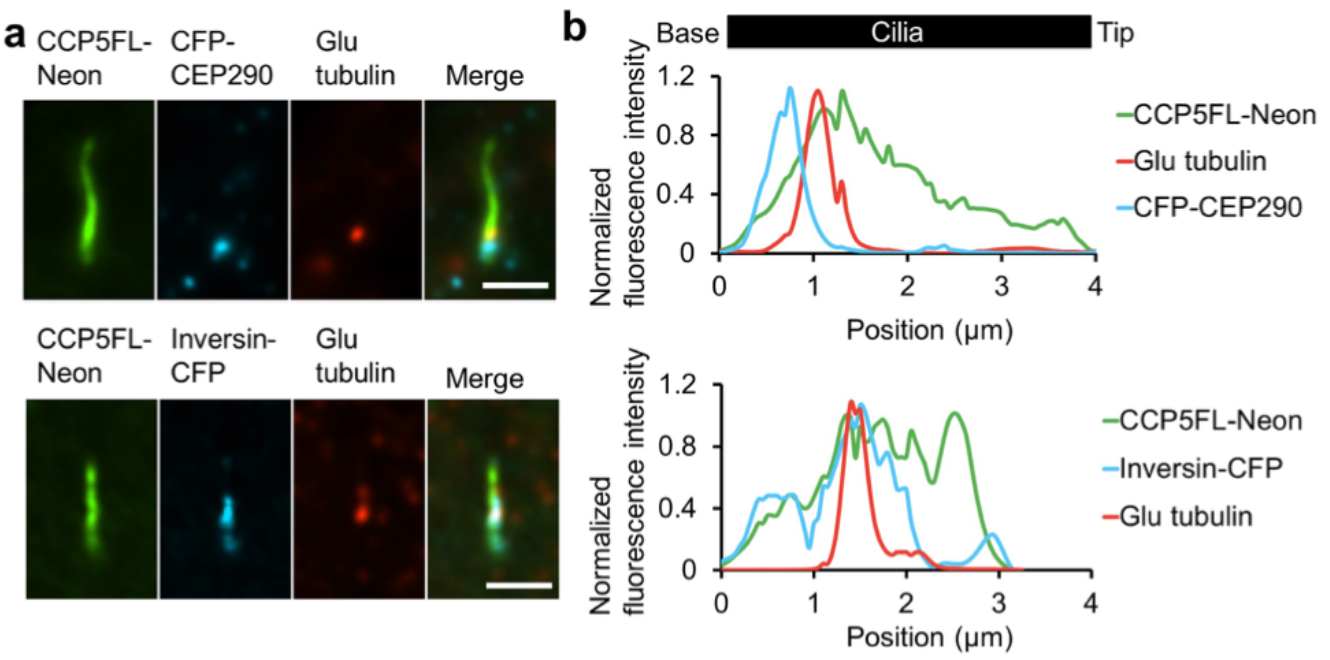
The distribution of residual glutamylated tubulin after axonemal deglutamylation. (**a**) Residual glutamylated tubulin localize at the inversin zone but not the transition zone after deglutamylation. NIH3T3 were transfected with CCP5FL-Neon and CFP-CEP290 or Incersin-CFP constructs for 24 h. Transfected cells at 80∼90% confluency were serum starved for 24 h and then immunostained with anti-Glutamylated tubulin antibody. Scale bar, 2 μm. (**b**) Linescan profiles of the indicated proteins from the base to the tip of cilia in **a**.

**Supplementary Figure 7.**
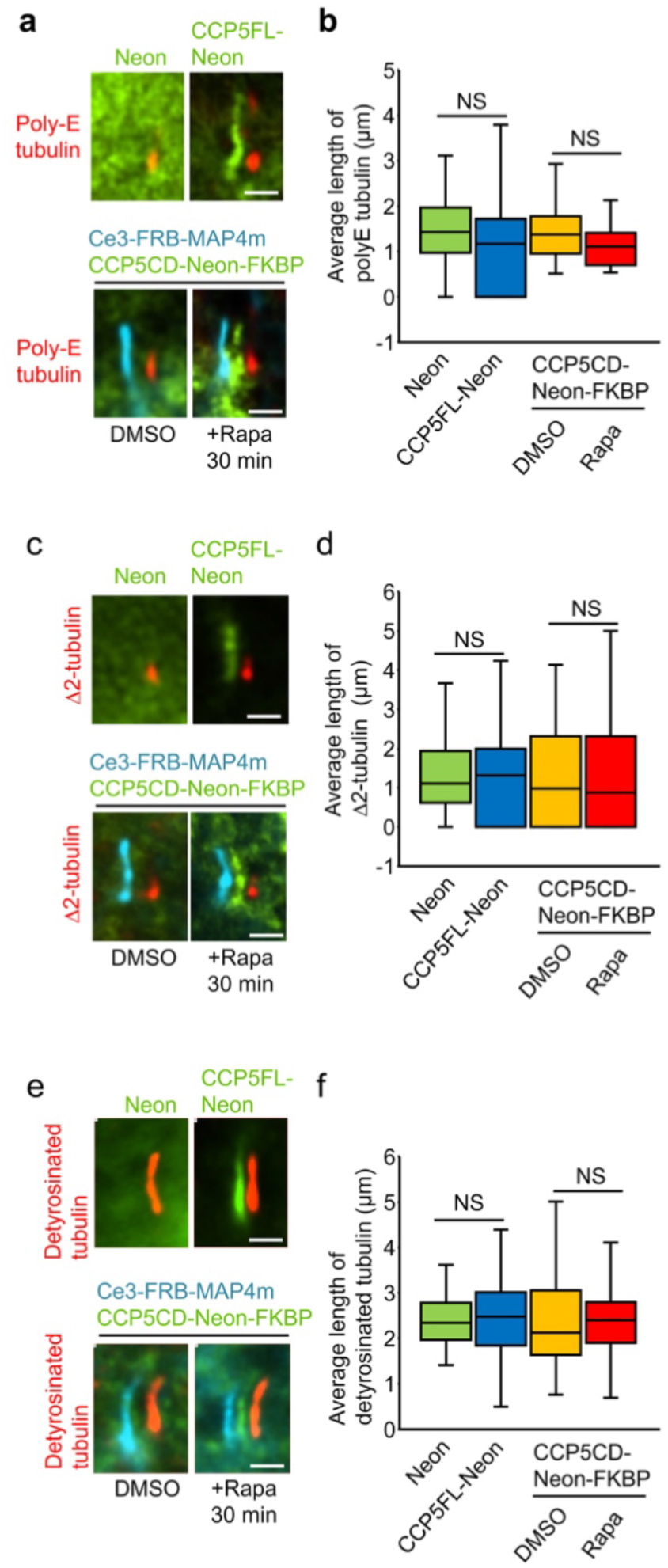
The enzyme activity of STRIP in poly-glutamate side chain shortening, Δ2-Δ3 tubulin conversion, and tubulin detyrosination. (**a,b**) CCP5 in cilia does not affect axonemal poly-glutamate (poly-E) (**a,b**), Δ2-tubulin (**c.d**), or axonemal detyrosination (**e, f.**). NIH3T3 cells were transfected with Neon, CCP5FL-Neon, or Cerulean3(Ce3)-FRB-MAP4m-P2A-CCP5CD-Neon-FKBP. Transfected cells at 80∼90% confluency were serum starved for 24 h, followed by a treatment of 0.1% DMSO or 100 nM rapamycin for 30 min. The level of ciliary poly-glutamate side chain, Δ2-tubulin, and tubulin detyrosination in cells expressing the indicated proteins upon 0.1% DMSO or 100 nM rapamycin treatment (Rapa) for 30 min was assessed by labeling with specific antibodies and were quantified. The images show shifted overlays of indicated proteins in cilia. Scale bar, 2 μm. (n = 169, 121, and 291 cells in the poly-E, Δ2-tubulin, and tubulin detyrosination experiments, respectively; three independent experiments). NS indicate no significant difference between the control and the indicated groups.

**Supplementary Figure 8.**
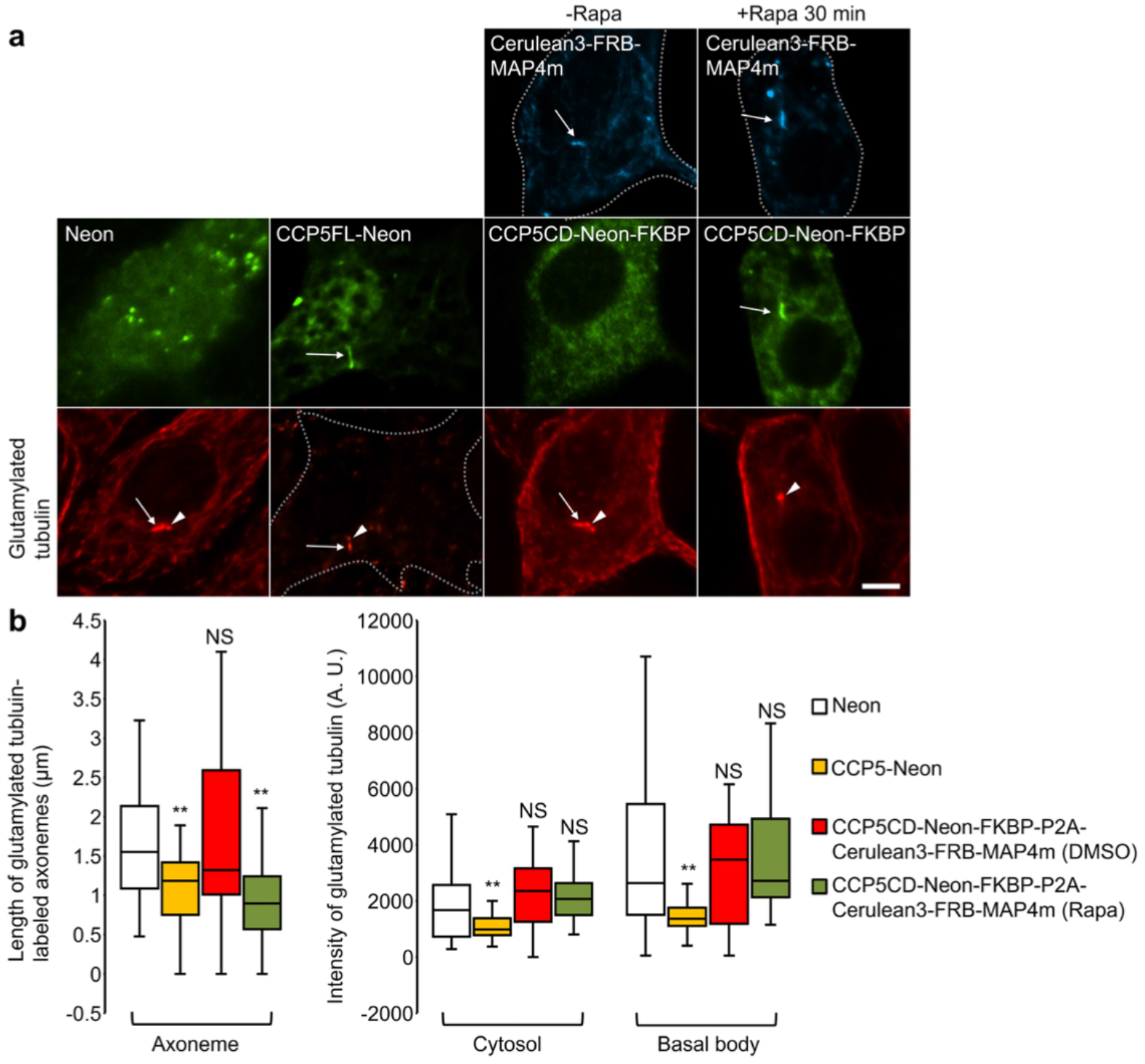
Specifically depleting axonemal glutamylation by our STRIP system. (**a**) CCP5FL-Neon but not CCP5CD-Neon-FKBP globally depletes glutamylation in NIH3T3 cells. Translocation of CCP5CD-Neon-FKBP onto the Cerulean3-FRB-MAP4m-labeled axoneme specifically strips the axonemal glutamylation without affecting the tubulin glutamylation in the cytosol and basal body. NIH3T3 cells were transfected with Neon, CCP5FL-Neon, or Cerulean3-FRB-MAP4m-P2A-CCP5CD-Neon-FKBP. Transfected cells at 80∼90% confluency were serum starved for 24 h and then treated with 0.1% DMSO (-Rapa) or 100 nM rapamycin (+Rapa) for 30 min. Subsequently, cells were incubated on ice for 1 h and immunostained with anti-glutamylated tubulin. Arrows and arrowheads mark the axoneme and basal body, respectively. Scale bars, 5 μm. (**b**) The length of glutamylated axoneme and the level of tubulin glutamylation in the cytosol and basal body in various conditions. (n = 71, 18, 25, 24, 206, 18, 34, 31, 197, 18, 29, 27 cells from left to right; 3-5 independent experiments). NS and ** represent no significant differece and *P* < 0.01, respectively, between the control (Neon) and the indicated conditions.

**Supplementary Figure 9.**
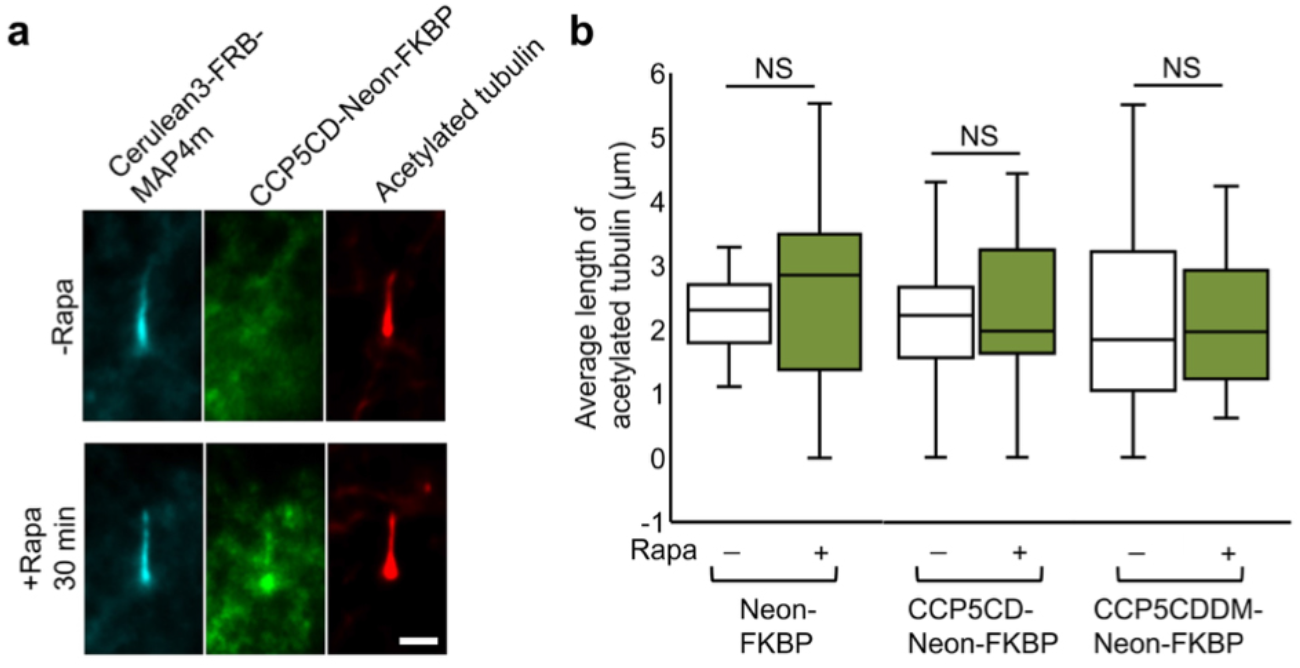
Rapid deglutamylation does not affect axonemal acetylation. (**a**) Translocation of CCP5CD onto the axoneme does not affect the level of acetylated tubulin. NIH3T3 cells were transfected with P2A-based constructs for co-expression of Cerulean3-FRB-MAP4m and Neon-FKBP-tagged proteins. Transfected cells at 80∼90% confluency were serum starved for 24 h and treated with or without 100 nM rapamycin for 30 min. Subsequently, cells were immunostained with anti-acetylated tubulin antibody. Scale bar, 2 μm. (**b**) Average length of acetylated tubulin in cells expressing the indicated proteins in the presence or absence of 100 nM rapamycin treatment for 30 min were quantified. (n = 13, 19, 20, 17, 19, 15 cells from left to right; three independent experiments). NS represents no significant difference between groups with or without rapamycin treatment.

**Supplementary Figure 10.**
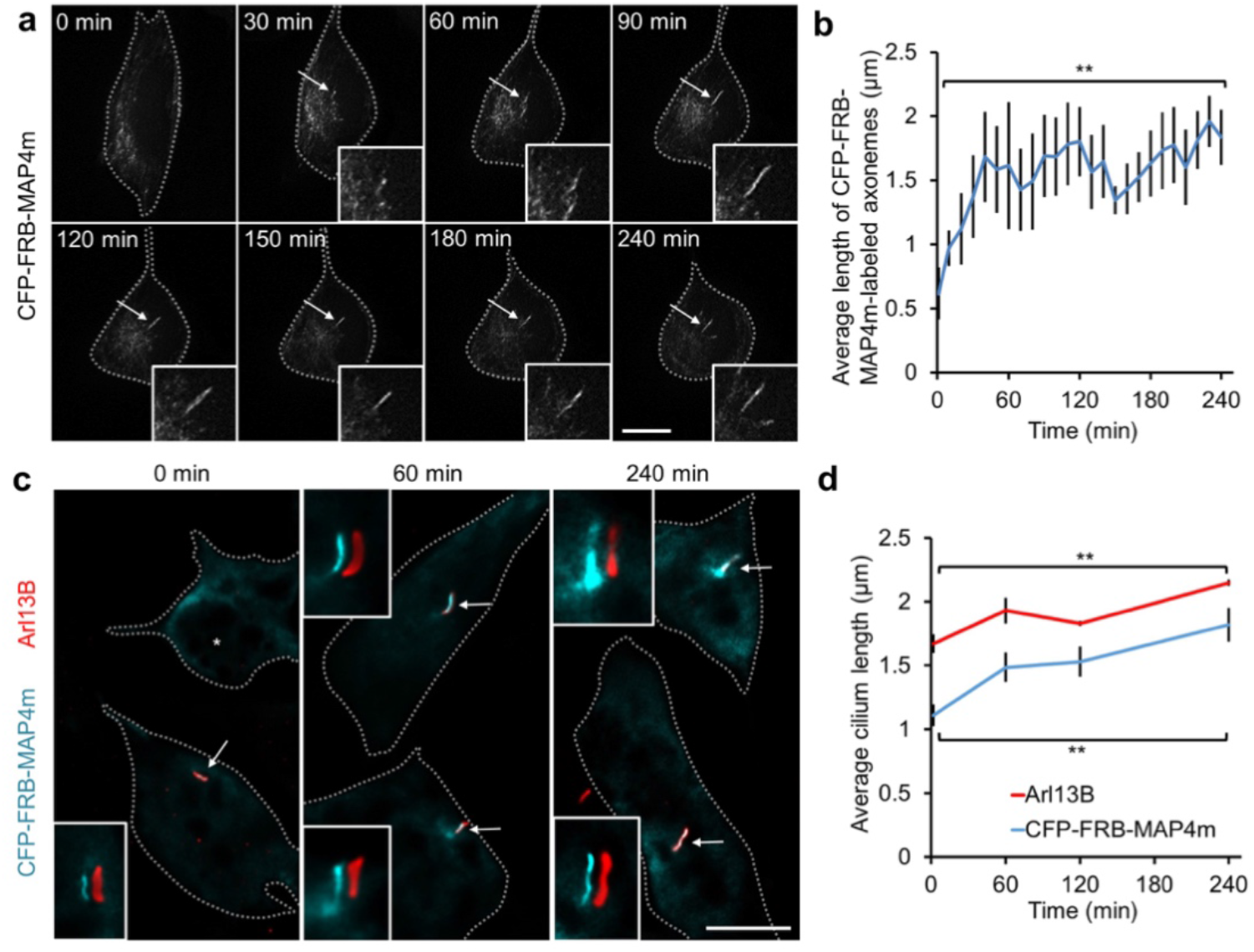
MAP4m serves as an axoneme targeting protein during ciliogenesis. (**a**) Time-lapse imaging of a NIH3T3 cell stably expressing CFP-FRB-MAP4m after serum starvation for the indicated times. Magnified views of the growing MAP4m-labeled axoneme (arrows) are shown in the insets. Scale bar, 10 μm. Also see Supplementary Movie 5. (**b**) Average length of the MAP4m-labeled axoneme after serum starvation for the indicated times was quantified. (n = 8 cells; two independent experiments). (**c**) The cilia in NIH3T3 cells stably expressing CFP-FRB-MAP4m (green) after serum starvation for the indicated times were immunostained with anti-Arl13B antibody (red). The insets show shifted overlays of MAP4m and Arl13B (arrows). A non-ciliated cell is marked by an asterisk. Scale bar, 10 μm. (d) Average length of the MAP4m-labeled axoneme and the Arl13B-labeled ciliary membrane after serum starvation for indicated times was quantified. (n≥ 180 cells; three independent experiments). Data represent the mean ± s.e.m. ** represent *P* < 0.01 between the cilium length before and after serum starvation for 240 min.

**Supplementary Figure 11.**
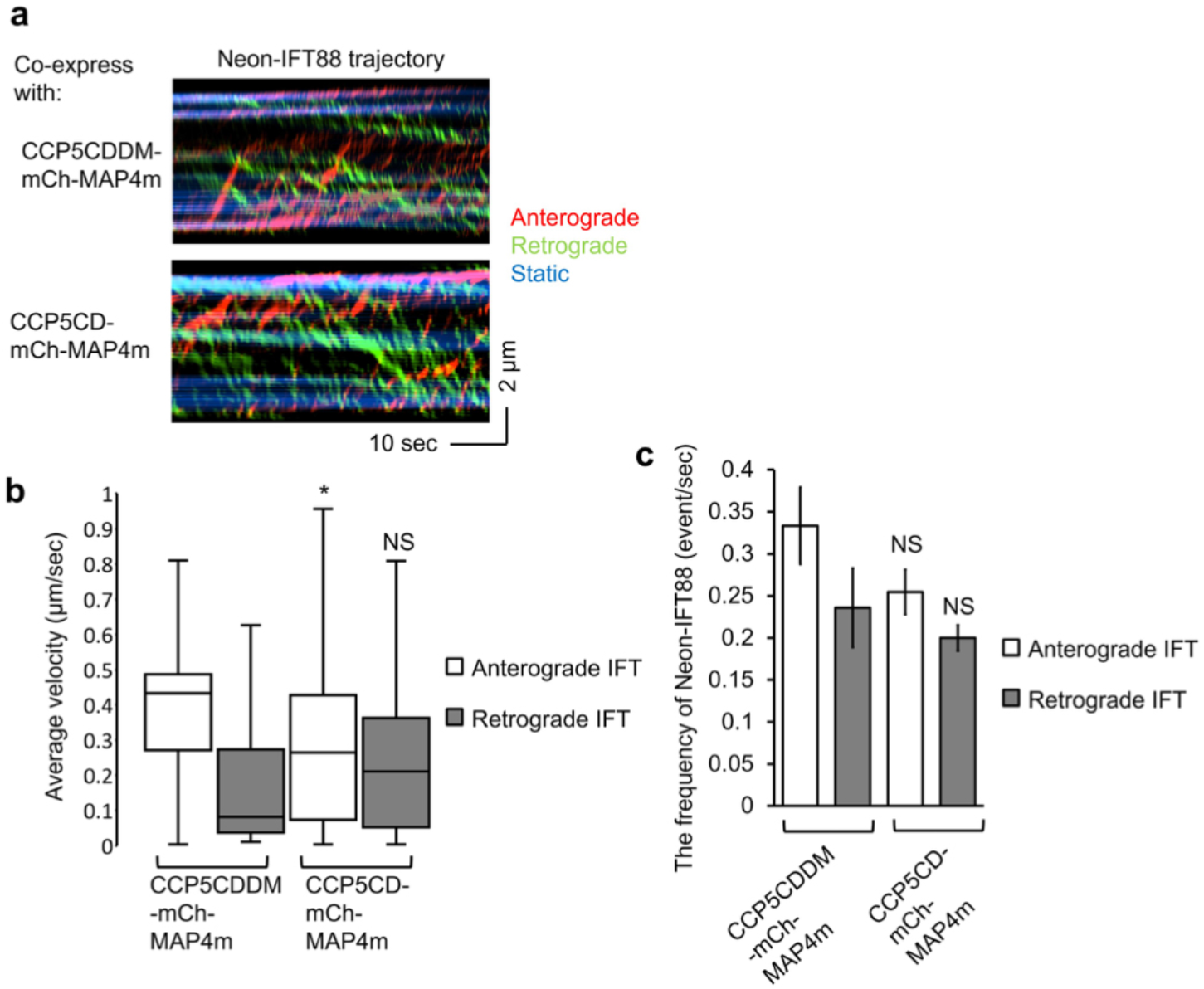
Long-term deglutamylation preferentially hampers anterograde IFT. (**a**) The Neon-IFT88 stable NIH3T3 cells were transfected with CCP5CD-mCherry-MAP4m or catalytically inactive CCP5CDDM-mCherry-MAP4m. Transfected cells at 80∼90% confluency were serum starved for 24 h. Representative kymographs of Neon-IFT88 in cells co-expressing CCP5CD-mCherry-MAP4m or CCP5CDDM-mCherry-MAP4m were generated from time-lapse imaging together with KymographClear. Red, green, and blue lines represent the trajectories of Neon-IFT88 particles in anterograde, retrograde directions and static Neon-IFT88, respectively. (**b**) The velocity of Neon-IFT88 in anterograde and retrograde direction was quantified according to the trajectories shown in kymogrpahs (See the method). (n = 353, and 261 neon-IFT88 particles in the CCP5CD-mCherry-MAP4m and CCP5CDDM-mCherry-MAP4m groups, respectively; three independent experiments). (**c**) The frequency of Neon-IFT88 particles moving along cilia expressing CCP5CD-mCherry-MAP4m or CCP5CDDM-mCherry-MAP4m (n = 13 and 11 cilia in the CCP5CD-mCherry-MAP4m and CCP5CDDM-mCherry-MAP4m groups, respectively; three independent experiments). Data represent the mean ± s.e.m. NS and * indicate no significant difference and *P* < 0.05, respectively, between the CCP5CDDM-mCherry-MAP4m and the CCP5CD-mCherry-MAP4m groups.

**Supplementary Figure 12.**
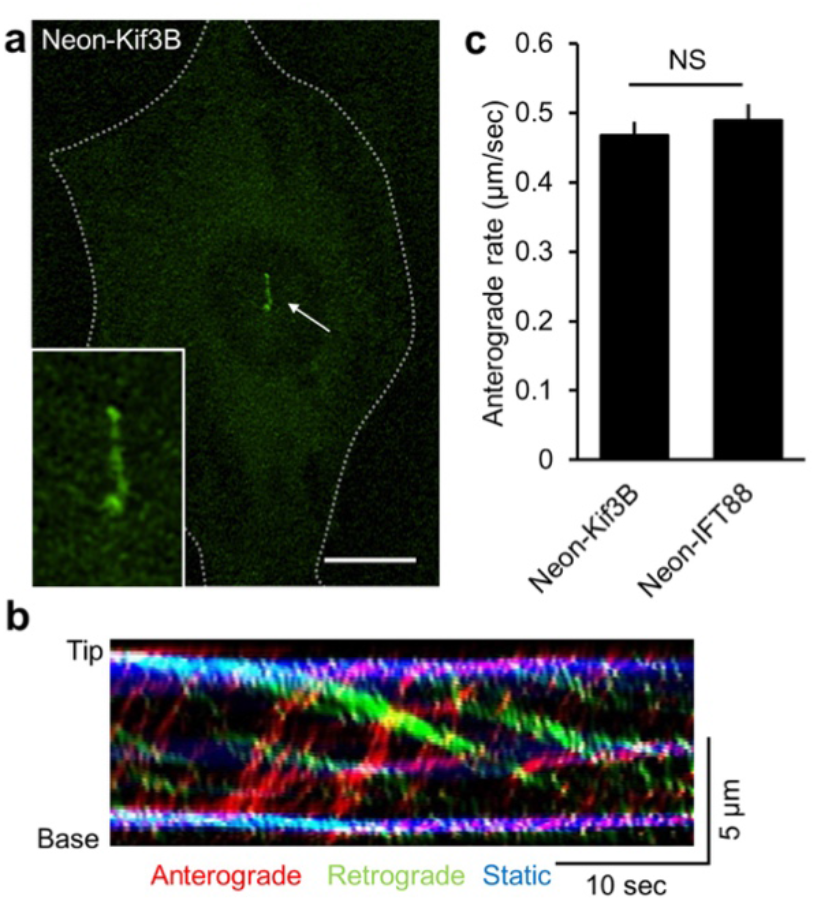
Neon-Kif3B localizes and moves along the axoneme. (**a**) The distribution of Neon-Kif3B in NIH3T3 cells are shown. Magnified view of the cilium expressing Neon-Kif3B (arrow) is shown in the inset. Scale bar, 10 μm. (**b**) A kymograph was generated from time-laspe imaging of cilium expressing Neon-Kif3B. The trajectories of Neon-Kif3B particles in anterograde and retrograde direction are highlighted by blue and red lines, respectively. (**c**) The velocity of Neon-Kif3B and Neon-IFT88 in anterograde direction. (n = 258, 158 IFT particles from left to right; four independent experiments. Data represent the mean ± s.e.m. NS represent no significant difference between the Neon-Kif3B and the Neon-IFT88.

**Supplementary Figure 13.**
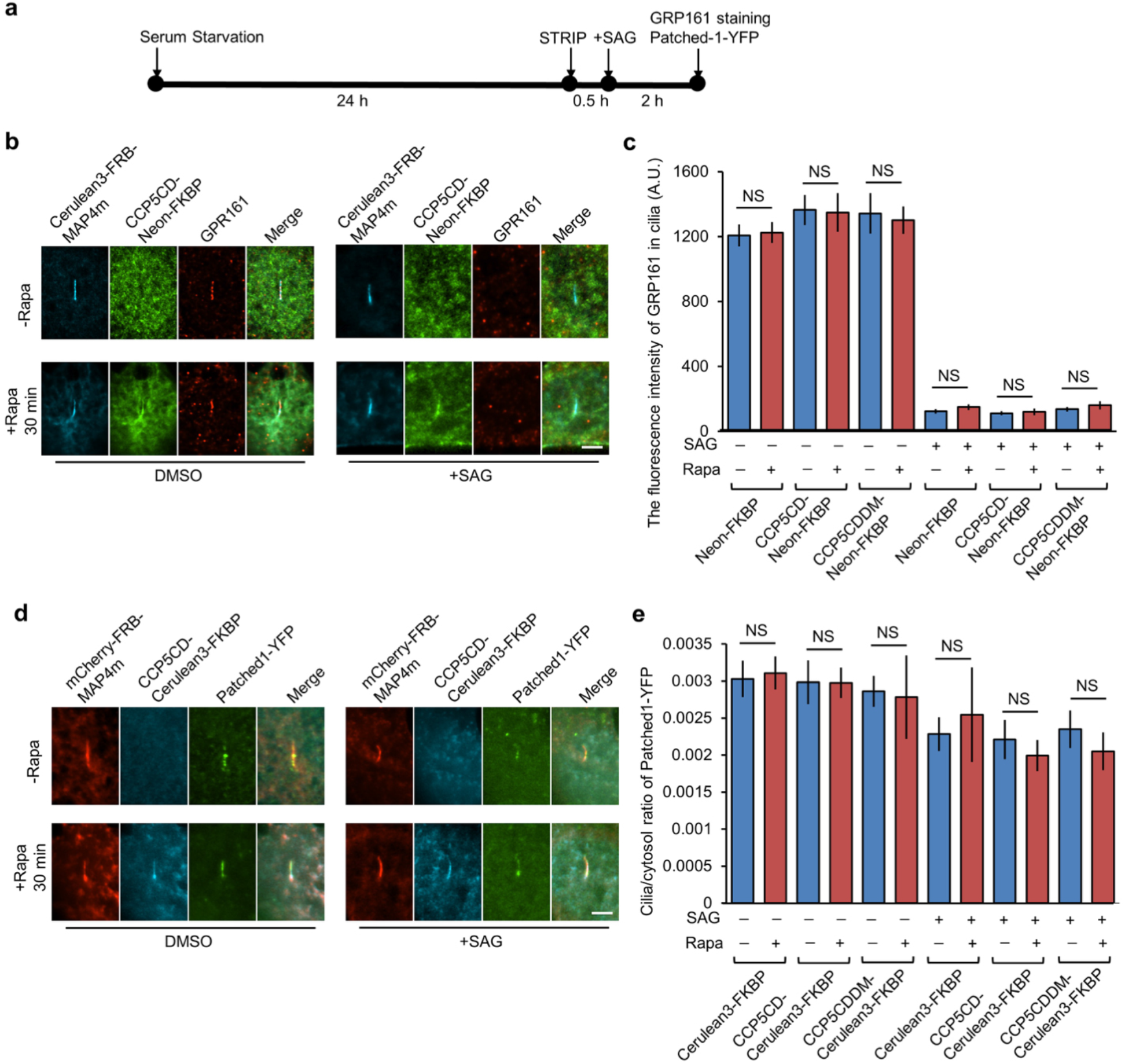
Axonemal deglutamylation does not affect the ciliary exit of GPR161 and Patched1. (**a**) The experimental procedure for STRIP and Hedgehog induction. Washout of rapamycin after 30 min STRIP operation did not affect protein dimerization in cells owing to the irreversible nature of the CID system. (**b**) STRIP-induced axonemal deglutamylation does not affect the ciliary exit of GPR161 upon stimulation of the SAG. NIH3T3 cells were transfected with P2A-based constructs for co-expression of Cerulean3-FRB-MAP4m and Neon-FKBP-tagged proteins. Transfected cells at 80∼90% confluency were serum starved for 24 hrous, followed by a treatment of 0.1% DMSO (-Rapa) or 100 nM rapamycin (+Rapa) for 30 min. Subsequently, cells were treated with 1 μM SAG for 2 h and immunostained with anti-GRP161 antibody. Sacle bar, 2 μm. (**c**) The fluorescence intensity of GRP161 in cilia was quantified and plotted (n = 318, 235, and 218 cilia in the Neon-FKBP, CCP5CD-Neon-FKBP, and CCP5CDDM-Neon-FKBP groups, respectively; 3-5 independent experiments). Data represent the mean ± s.e.m. NS represents no significant difference between with and without rapamycin treatment group. (**d**) STRIP-induced axonemal deglutamylation does not affect the ciliary exit of Patched1-YFP upon stimulation of the SAG. NIH3T3 cells were transfected with Patched1-YFP and P2A-based constructs for co-expression of mCherry-FRB-MAP4m and Cerulean3-FKBP-tagged proteins. Transfected cells at 80∼90% confluency were serum starved for 24 h and then treated with 0.1% DMSO (-Rapa) or 100 nM rapamycin (+Rapa) for 30 min. Subsequently, cells were treated with 1 μM SAG for 2 h. Scale bar, 2 μm. (**e**) The ratio of Patched1-YFP intensity in cilia and cytosol was quantified and plotted. (n = 201, 210, and 132 cilia in the Cerulean3-FKBP, CCP5CD-Cerulean3-FKBP, and CCP5CDDM-Cerulean3-FKBP groups, respectively; three independent experiments).

**Supplementary Figure 14.**
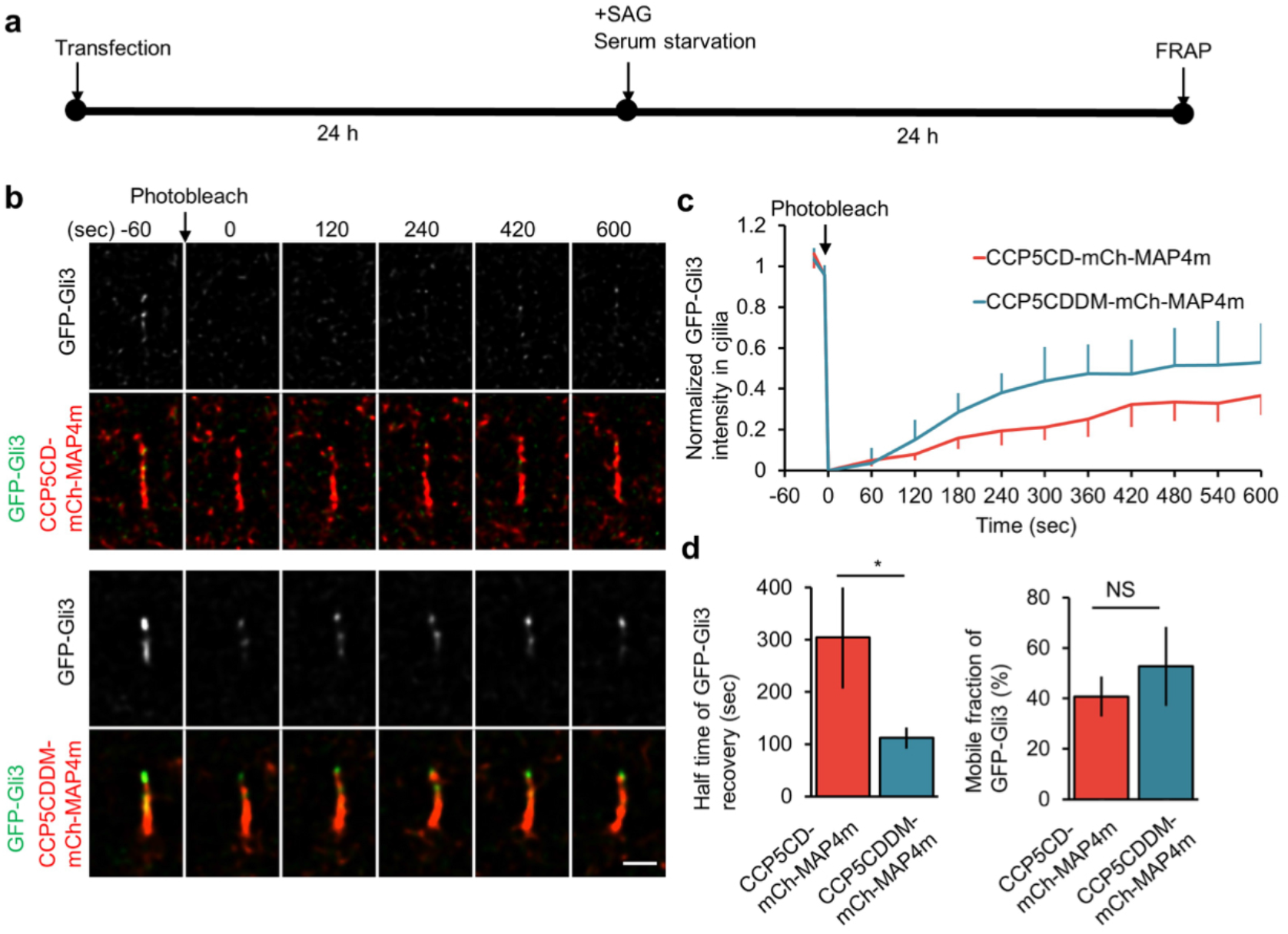
Axonemal deglutamylation hampers ciliary entry of Gli3 upon SAG stimulation. (**a**) The experimental procedures in **b**. NIH3T3 cells were transfected with GFP-Gli3 and CCP5CD-mCherry-MAP4m or CCP5CDDM-mCherry-MAP4m for 24 h. Transfected cells at 80∼90% confluency were treated with 200 nM SAG in serum free medium for 24 h and then analyzed by FRAP. (**b**) The GFP-Gli3 in cells expressing CCP5CD-mCherry-MAP4m or catalytically inactive CCP5CDDM-mCherry-MAP4m was photobleached and allowed for recovery for 10 min. Scale bar, 1 μm. (**c**) The fluorescence recovery of GFP-Gli3 in cilia expressing CCP5CD-mCherry-MAP4m or CCP5CDDM-mCherry-MAP4m was measured and plotted. (**d**) The recovery rate (left) and mobile fraction (right) of GFP-Gli3 in cilia were measured and plotted. Data represent the mean ± s.e.m. (n = 5, and 8 cilia for the CCP5CD-mCherry-MAP4m and CCP5CDDM-mCherry-MAP4m groups, respectively; three independent experiments). NS and * represent no significant difference and *P* < 0.05, respectively, between the CCP5CDDM-mCherry-MAP4m and the CCP5CD-mCherry-MAP4m groups.

